# Population structure and history of *Mycobacterium bovis* European 3 clonal complex reveal transmission across ecological corridors of unrecognised importance in Portugal

**DOI:** 10.1101/2023.11.01.564909

**Authors:** André C. Pereira, José Lourenço, Gonçalo Themudo, Ana Botelho, Mónica V. Cunha

## Abstract

*Mycobacterium bovis* causes animal tuberculosis in livestock and wildlife, with an impact on animal health and production, wildlife management, and public health. In this work, we sampled a multi-host tuberculosis community from the official hotspot risk area of Portugal over 16 years, generating the largest available dataset in the country. Using phylogenetic and ecological modelling, we aimed to reconstruct the history of circulating lineages across the livestock-wildlife interface to inform intervention and the implementation of genomic surveillance within the official eradication plan. We find evidence for the co-circulation of *M. bovis* European 1 (Eu1), Eu2, and Eu3 clonal complexes, with Eu3 providing sufficient temporal signal for further phylogenetic investigation. The Eu3 most recent common ancestor (bovine) was dated in the 90’s, subsequently transitioning to wildlife (red deer and wild boar). Isolate clustering based on sample metadata was used to inform phylogenetic inference, unravelling frequent transmission between two clusters that represent an ecological corridor of previously unrecognised importance in Portugal. The latter was associated with transmission at the livestock-wildlife interface towards locations with higher temperature and precipitation, lower agriculture and road density, and lower host densities. This is the first analysis of *M. bovis* Eu3 complex in Iberia, shedding light on background ecological factors underlying long-term transmission, and informing where efforts could be focused within the larger hotspot risk area of Portugal.

**IMPORTANCE:** Efforts to strengthen surveillance and control of animal tuberculosis (TB) are ongoing worlwide. Here, we developed an eco-phylodynamic framework based on discrete phylogenetic approaches informed by *M. bovis* whole genome sequence data representing a multi-host transmission system at the livestock-wildlife interface, within a rich ecological landscape in Portugal, to understand transmission processes and translate this knowledge into disease management benefits.

We find evidence for the co-circulation of several *M. bovis* clades, with frequent transmission of the Eu3 lineage among cattle and wildlife populations. Most transition events between different ecological settings took place toward host, climate and land use gradients, underscoring animal TB expansion and a potential corridor of unrecognised importance for *M. bovis* maintenance.

Results stress that animal TB is an established wildlife disease without ecological barriers, showing that control measures in place are insufficient to prevent long-distance transmission and spillover across multi-host communities, demanding new interventions targeting livestock-wildlife interactions.

## INTRODUCTION

*Mycobacterium bovis* is the causative agent of animal tuberculosis (TB) in livestock and wildlife, retaining zoonotic potential [1]. The main affected livestock worldwide is *Bos taurus* (bovine), and in the Iberian Peninsula, the main affected wildlife species are *Sus scrofa* (wild boar) and *Cervus elaphus* (red deer) [2, 3, 4]. The livestock-wildlife interface is considered of key importance in the dissemination of *M. bovis* by both direct and indirect transmission routes, which are highly dependent on husbandry systems and host activity aggregation points [5, 6, 7].

In Portugal, there is a national bovine TB program aiming control towards eradication [8]. It is based on three pillars: (i) detection and compulsory slaughter of animal reactors to the single intradermal comparative cervical tuberculin test; (ii) routine surveillance of carcasses at slaughterhouses; and (iii) pre-movement testing [8]. The official wildlife surveillance program targets a hotspot risk area related to the synanthropy of big game species due to natural conditions and artificial management, in east-central and eastern-south mainland Portugal [5, 6, 9].

*M. bovis* epidemiological surveillance in Portugal has mostly been based on classic molecular characterization, including spoligotyping and MIRU-VNTR [6, 10, 11]. Efforts remain focused on the three main reservoirs of TB within the hotspot risk area [6, 10, 11]. Classic molecular approaches analyse only <1% of the genome and are not sufficiently discriminatory to accurately assess transmission chains. Outputs are particularly compromised by homoplasy and are insufficient to gain insights into the roles exerted by different species in the multi-host system [12]. Whole-genome sequencing (WGS) approaches can overcome these limitations, using single nucleotide polymorphisms (SNPs) as reliable phylogenomic markers, identifying genetic population structures while linking to co-variables of interest, thus unraveling transmission drivers and routes [13, 14, 15, 16]. In recent, first of a kind pilot studies applying WGS, we have identified transmission between livestock (bovine) and wild ungulates (red deer, wild boar) in the Castelo Branco and Portalegre districts within the hotspot risk area of mainland Portugal [11, 16]. Global studies have identified five main clonal complexes, with three predominant in Europe: Eu1 is globally distributed, with an origin associated with the British Isles [17]; Eu2 (lineage La1.7.1) is predominant in the Iberian Peninsula [17]; Eu3 was recently shown to be predominant in Western Europe and East-Africa [18]. In this study, under a new sampling and WGS effort over 16 years (2002-2018), we substantially increase the number of existing *M. bovis* full genomes (from 44 to 170) from within the hotspot risk area in mainland Portugal. Using phylogenetic and ecological modelling approaches informed by WGS and sample metadata, we recover and describe local transmission chains, provide robust estimations of several evolutionary parameters, and uncover an ecological corridor of previously unrecognised importance in mainland Portugal.

## RESULTS

### Population Structure

The novel dataset reflects the local surveillance history of TB, which is characterised by: low sampling before 2006 due to insufficient surveillance (**Figure 1A**); an increase in wildlife isolates after 2011 due to the creation of the official hotspot risk area related to wildlife game species (**Figure 1A**); and by an overall higher number of isolates recovered from Castelo Branco due to a higher hunting activity in that area (**Figure 1B**). The phylogenetic distribution of SNPs grouped isolates into clades 1 to 10 (**Figure 1C**). Clades 1 to 7 were related to Eu2, clade 8 to Eu1, clade 9 to Unk7 (unknown clonal complex 7, also known as lineage 1.8.2), and clade 10 to Eu3 (**Figure 1C, Supplementary Table T1S2**). All major clades included strains from the three host species and both districts, suggesting mixing in transmission sources and routes.

**Figure 1.**
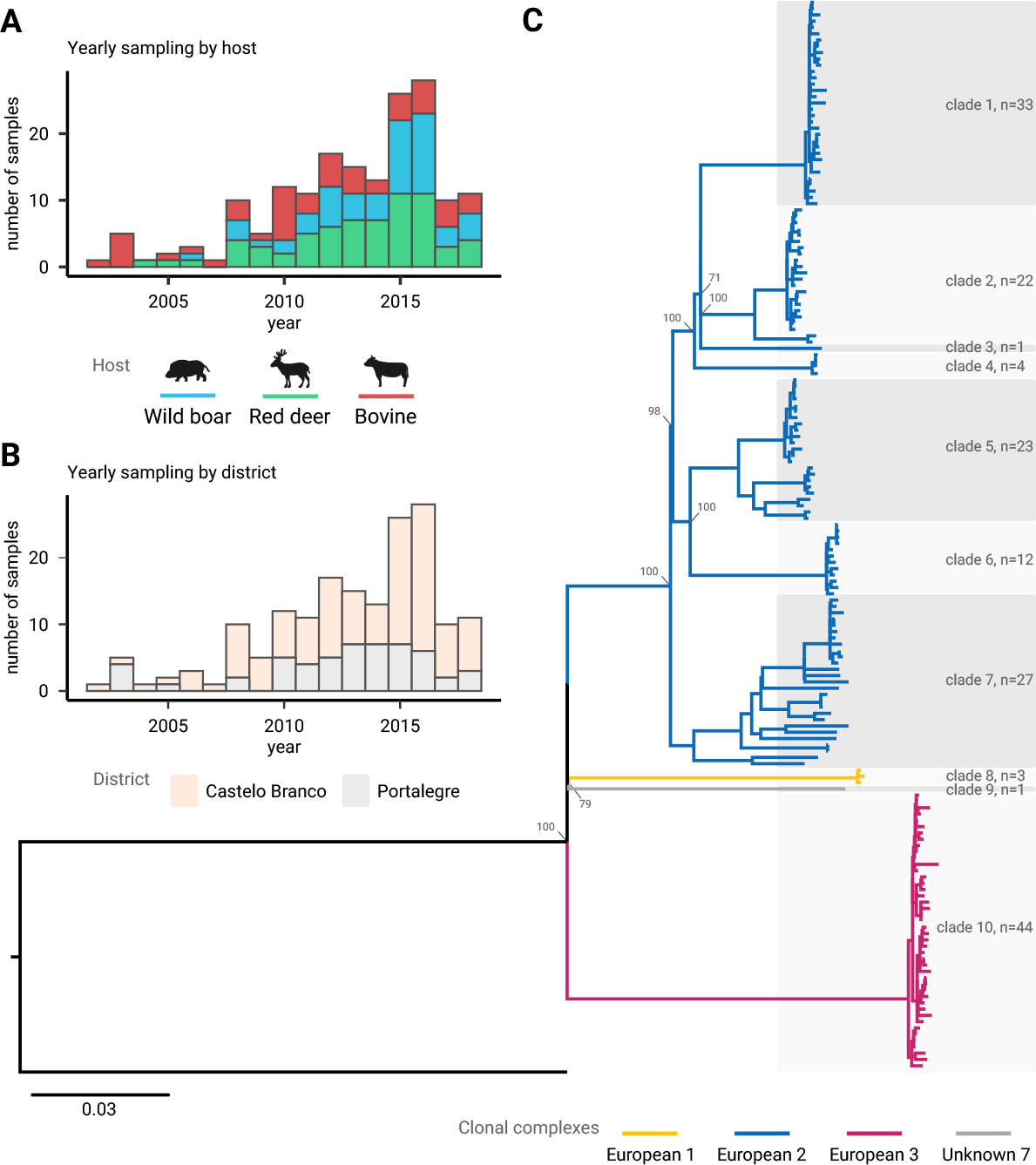
*M. bovis* sampling according to host and district and maximum likelihood phylogenetic tree. Total number of samples per year in the dataset according to (A) host (wild boar in blue, red deer in green, bovine in red) and (B) district (Castelo Branco in orange, Portalegre in grey). (C) Maximum likelihood SNP-based phylogenetic tree showing clades 1 to 10 (with sampling sizes), and colour coded major clades corresponding to clonal complexes: European 1 (yellow, including clade 8), European 2 (blue, including clades 1-7), European 3 (magenta, including clade 10) and Unknown 7 (grey, including clade 9).

### Transmission Mapping

Local transmission networks based on the SNP alignment and available metadata identified several transmission chains (**Figure 2**). Fifteen chains were identified with 2 to 5 isolates per chain (25 events). In Castelo Branco district, there were nine transmission chains, with two at the wildlife-livestock interface, one in livestock, six in wildlife including three within the same hunting area. In Portalegre district, there were five transmission chains, with two at the wildlife-livestock interface, two in livestock involving animals from within the same herd, and one exclusive to wildlife. No inter-district transmission events were found. Only three in twenty five inferred individual transmission events had zero SNP differences and all occurred within the same host species (bovine). Nevertheless, strains with less than six SNP differences were also detected within a time interval of six to eleven years even within the same hunting area.

**Figure 2.**
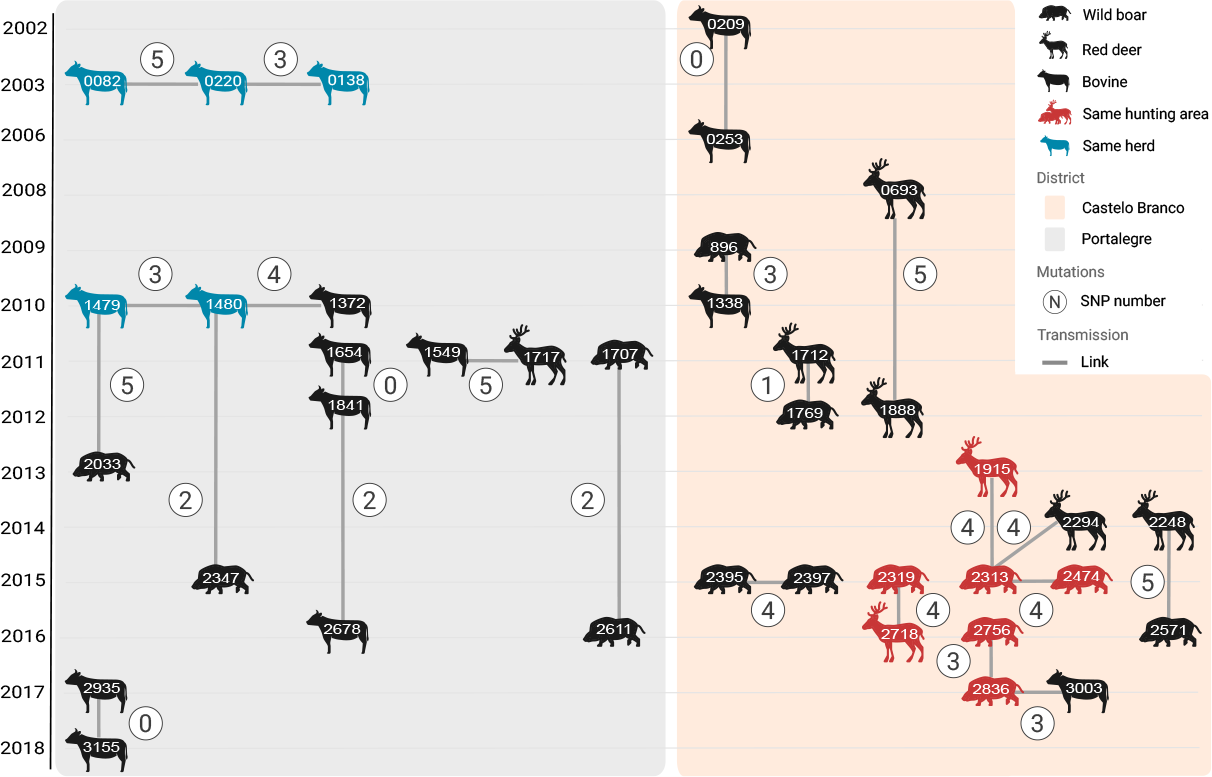
Inferred recent *M. bovis* transmission network. Diagram of the recovered transmission events according to host species (symbols), individuals (ID number), location (background shading), time (scale on the left), and SNP difference (circles with numbers). Lines mark transmission links between individuals. Individuals within the same hunting area are in red (wildlife) and those within the same herd are in blue (bovine).

### Phylogenomics

Root-to-tip and date-randomization analyses confirmed that Clade 10 (Eu3, **Figure 1C**) presented sufficient temporal signal for further phylogenomic analyses (**Supplementary Figure T1S2**). The clade had 44 isolates (25% of isolates) from bovine (n=13), red deer (n=16), and wild boar (n=15), from Castelo Branco (n=18) and Portalegre (n=26), sampled in 2003 and 2010-2018. TIM2 with invariant sites and a four categories discrete gamma-distribution (TIM2+I+G) was the best-fitting nucleotide substitution model (**Supplementary Table T1S4**).

A time-based phylogeny indicated the relaxed exponential clock with a Bayesian Skyline population as the best-fitting model (**Supplementary Table T1S5**). The mean clock rate of Eu3 was estimated to be 2.2x10^-4^ (95% highest probability density, HPD, [8.9x10^-5^– 3.6x10^-4^]) substitutions per site per year, corresponding to a mean evolutionary rate of 0.2 (95% HPD [0.1–0.4]) substitutions per genome per year. The median tMRCA was estimated 28 years ago (95% HPD [17–53]), corresponding to 1991 (95% HPD [1965– 2001]) (**Figure 3A**). A lineage through time analysis (**Figure 3B**) revealed that after the tMRCA, an initial increase in strain diversity occurred, with particular growth around the turn of the century and eventual plateauing around 2010. A coalescent Bayesian Skyline analysis estimating the effective population size (Ne) through time, further supported these temporal observations (**Figure 3C**).

**Figure 3.**
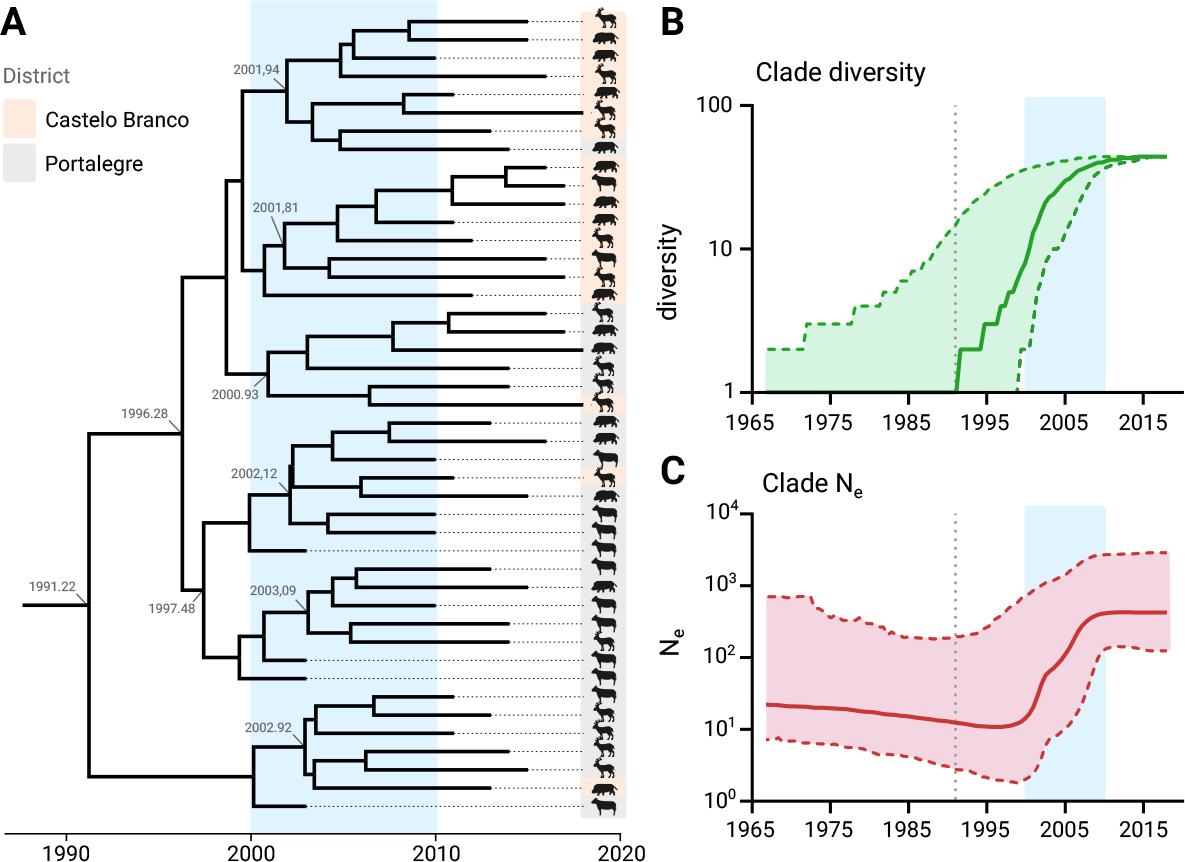
European 3 clonal complex ancestry, diversification, and effective population size. (A) Time-calibrated maximum clade credibility tree. Sampled individuals are presented by corresponding animal symbols and locations by shaded colours (orange for Castelo Branco, grey for Portalegre) on the right side. Numbers in grey along the tree show the branching timings of subclades. (B) Lineage-through-time analysis showing clade diversity in time (green), where the mean is represented by a full line and the 95% CI by a shaded area. (B) Lineage-through-time analysis showing clade diversity in time (green), where the median is represented by a full line and the 95% CI by a shaded area. (C) Bayesian skyline reconstruction shows effective population size in time (red), where the median is represented by a full line and the 95% CI by a shaded area. In both (B,C) panels the vertical dotted line marks the clade’s tMRCA, and in all panels, the blue shaded area highlights a period of particular growth.

### Ancestral State Host Reconstruction

We estimated the internal node probability of association with each host species. The symmetric model showed the best fitting (**Supplementary Table T1S6, Figure T1S3**). A slightly stronger signal was inferred for host transitions between both wildlife hosts (PP=0.87) compared to transitions between livestock and wildlife (bovine-red deer, PP=0.79; bovine-wild boar, PP=0.77) (**Figure 4A**). The asymmetric model, while less supported, also estimated reasonably high transitions (PP>0.5) between pairs of hosts, except for transitions between red deer and bovine (**Figure 4B**). The MRCA was associated with bovine, subsequently bifurcating into two subclades (clades 1, 2) (**Figure 4C**). Clade 1 was characterised by persistence in bovine and host transition events towards wildlife species, mostly within Portalegre, and clade 2 was characterised by persistence in wildlife, with later host transitions back to bovine, mostly within Castelo Branco.

**Figure 4.**
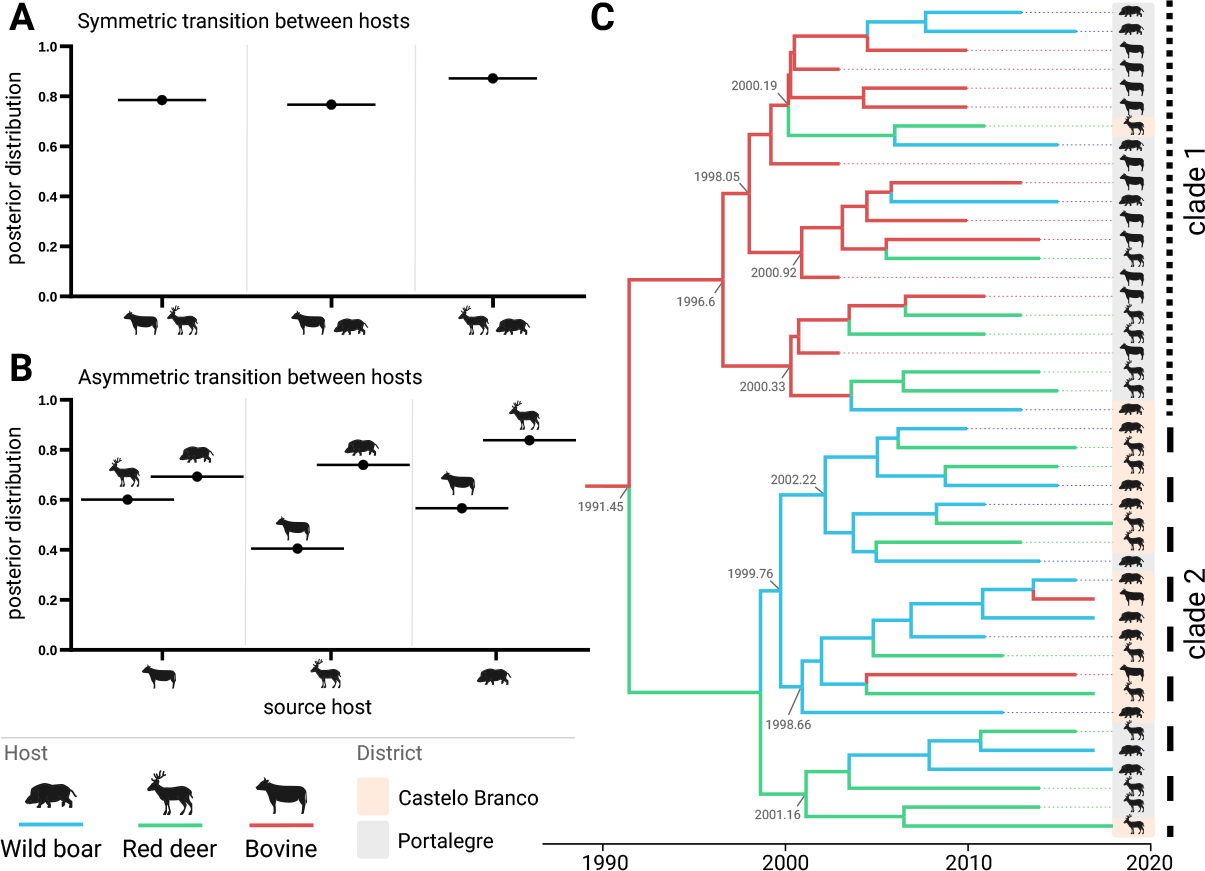
European 3 clonal complex ancestral state host reconstruction. (A) Host state posterior probabilities under a model of symmetric host species transitions, with source/sink hosts on the x-axis. (B) Host state posterior probabilities under a model of asymmetric host species transitions, with source host on the x-axis and sink host on the data points. (C) Maximum credibility tree estimated under a model of symmetric host species transitions. The two major clades 1 and 2 are highlighted on the right. Numbers in grey along the tree show the branching timings of subclades. In panels (A-C) host species are represented by animal symbols as well as colours (wild boar in blue, red deer in green, bovine in red), while location is represented by shaded background (orange for Castelo Branco and grey for Portalegre).

### Phylogeography Using Spatial Data

We next estimated the internal node probability relating to the sample’s geographic location. When comparing all phylogeographic models, using districts as grouping levels, applying a symmetric coalescent model was the most supported approach (**Supplementary Table T1S6**). The probability of transitions between both districts was close to 1.00 (**Supplementary Figure T1S3**). The results of all unsupported models are included in **Supplementary Text 1**.

The MRCA was associated with Portalegre district (municipality Castelo de Vide) with subsequent geographic events towards Castelo Branco district (**Figure 5A**). Similarly to the inferred timings of major growth in diversity and effective population size (**Figure 3BC**), a boost in inter-municipality transitions occurred after the turn of the XXI century. Estimated transitions between municipalities were recovered from the tree internal nodes, showing five municipalities dominating spatial transitions in time, namely Nisa (Portalegre district), Castelo de Vide (Portalegre), Portalegre (Portalegre), Idanha-a-Nova (Castelo Branco), and Castelo Branco (Castelo Branco) (**Figure 5B**). Notably, these five municipalities have higher wildlife animal density (**Figure 5C**). Castelo de Vide in the district of Portalegre was the municipality with most transitions, both intra- and inter-municipality.

**Figure 5.**
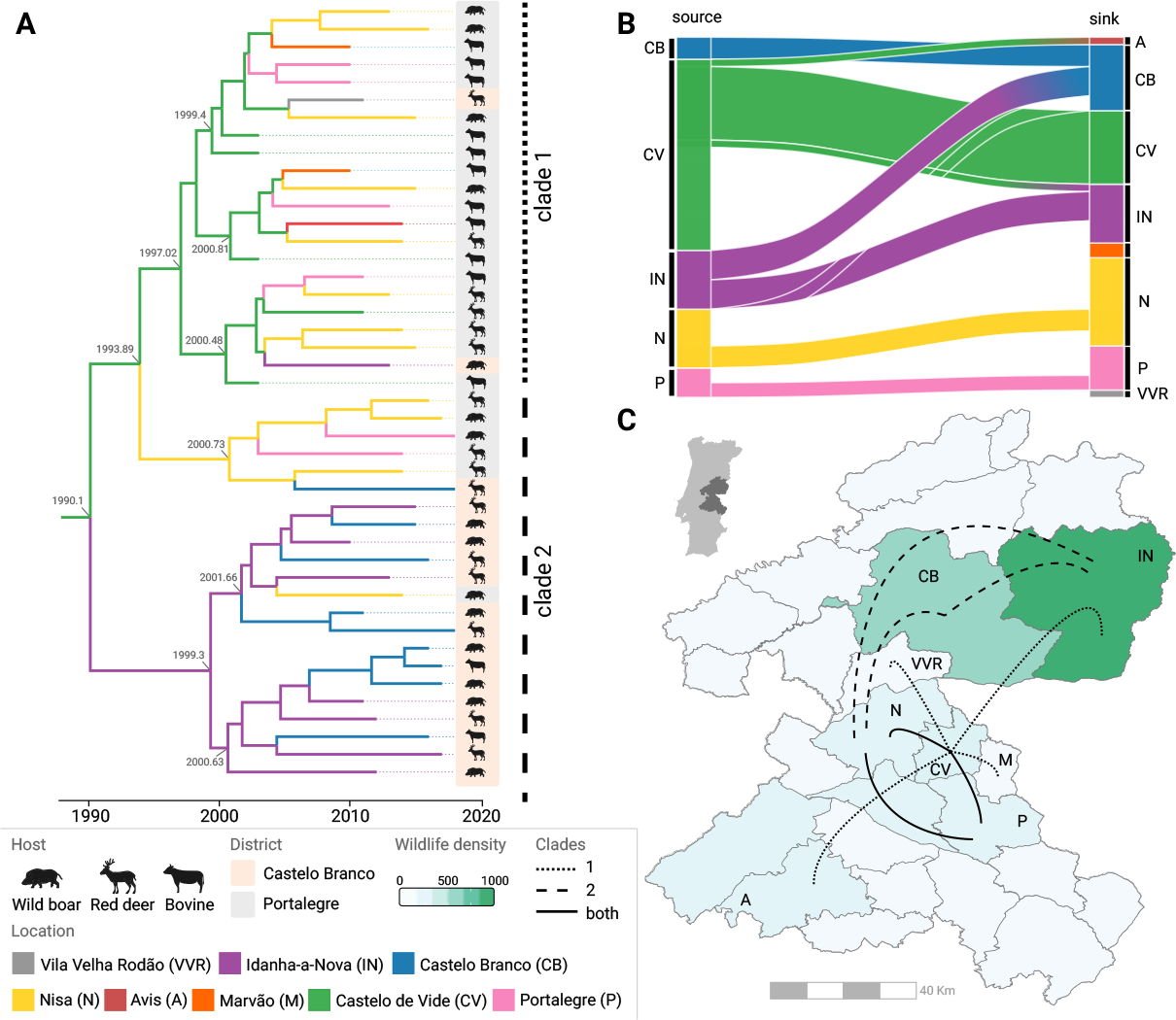
Phylogeography of the European 3 clonal complex using spatial boundaries and coordinates. (A) Maximum credibility tree estimated using a model of asymmetric municipality transitions. The two major clades 1 and 2 are highlighted on the right. Numbers in grey along the tree show the branching timings of subclades. Animal symbols are used to represent species and the shaded backgrounds highlight the district of sampling (orange for Castelo Branco, grey for Portalegre). (B) Alluvial plot denoting both intra- and inter-municipality transitions (recovered from inferences on internal nodes). In panels (A-B) colours are used to represent municipalities. Note that the names Portalegre and Castelo Branco are used both for districts and the respective main municipalities of those districts. (C) Spatial representation of inter-municipality transitions for clade 1 (dotted lines), clade 2 (shaded line), or both (full lines) on the background of wildlife density (green colour scale).

### Phylogeography Using Ecological Clustering Data

Finally, we estimated the internal node probability relating to the sample ecological cluster. When comparing all phylogeographic models, assuming the existence of four ecological clusters was the most supported approach (independently of the symmetry assumption used) (**Supplementary Table T1S6**).

The spatial distribution of the four ecological clusters showed geographical consistency across the study area (**Figure 6A**). Together, the first two dimensions of the MFA represented 61.6% of the total variance among the 170 samples (**Figure 6B**), revealing inherent correlative data structures. For example, while there was a positive correlation between the densities of red deer and mufflon, and independently between wild boar and bovine, these two groups of animals were negatively correlated. We mapped each sample and corresponding ecological cluster to the two first dimensions of the MFA (**Figure 6C**), which further revealed inherent data structures of interest (**Figure T2S8**). Ecological cluster 1 (n=94) was characterised by a combination of high mufflon and red deer densities together with the highest mean annual temperature (**Figure 6BC**). On the other hand, ecological cluster 2 (n=11) was associated with high agriculture and low forest land coverage, high bovine density and road density, and low mean annual temperature. Ecological cluster 3 (n=35) was not represented among the Eu3 samples, but among the remaining samples it appeared as an outlier of ecological cluster 1 with a much lower mean annual temperature (**Figure T2S8**). Finally, ecological cluster 4 (n=30) was linked to intermediate levels of several ecological variables and particularly low density of fallow deer and bovine. A full visual description of the ecological gradients across samples and clusters is in **Figure T2S8**.

**Figure 6.**
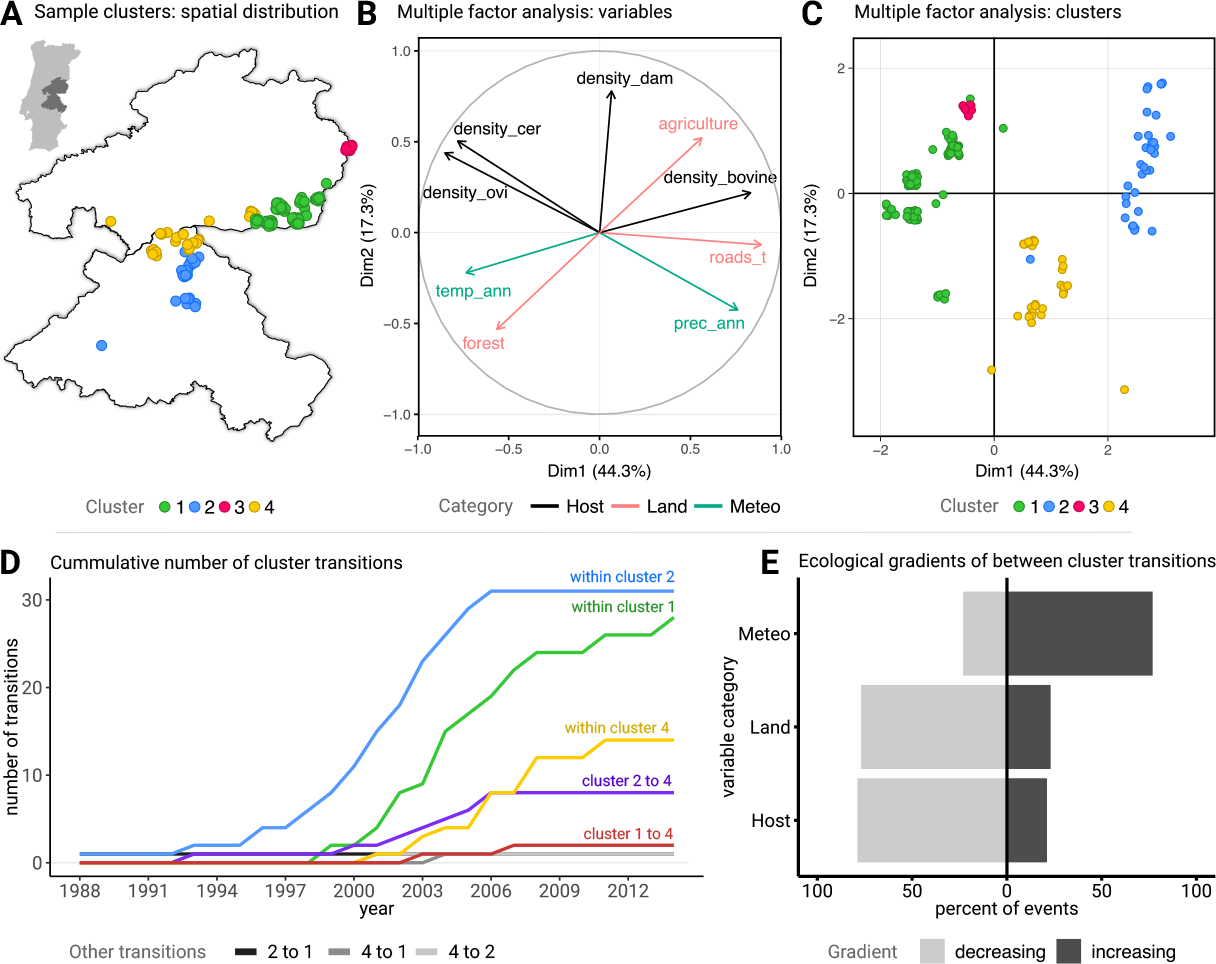
*M. bovis* ecological clustering and European 3 clonal complex cluster transitions. (A) Spatial distribution of ecological clusters (1 to 4, in different colours) where each point is an *M. bovis* isolate. (B) Multiple factor analysis (MFA) presenting dimensions 1 and 2. Shown are the top nine variables that most contributed to dimensions 1 and 2. Variables and their axes are coloured according to the variable category they belong to (black for those related to Host, pink when related to Land, and marine when related to Meteorology [Meteo]). For the definition of variable categories as for visual output including all variables, see Supplementary Text 2. (C) Mapping of each *M. bovis* isolate in MFA dimensions 1 and 2 space, coloured according to the ecological cluster. (D) Cumulative number of intra- and inter-cluster inferred transitions with time. (E) Inferred between cluster transitions are summarised depending on whether the mean value of ecological variables associated with clusters increased or decreased with the transition event. The decreasing label in the panel legend means that the cluster transition occurred within a negative ecological gradient in which the values of the variables included in the category (Host, Land, or Meteo) decrease (and vice-versa for the increasing label). In panels (D-E), only transitions related to the European 3 clonal complex are presented, for which we implemented phylodynamic inference.

Regarding Eu3 samples alone, phylogeographic transition probabilities between and within ecological clusters were estimated. The symmetric assumption showed high support for transitions between ecological clusters 1 and 4 (PP=0.89) and between ecological clusters 2 and 4 (PP=1.00). This was consistent with the inferred transition probabilities allowing asymmetry, where transitions from ecological cluster 1 to 4 (PP=0.84), from 2 to 1 (PP=0.66), and from 2 to 4 (PP=1.00) had high support. In general, the phylogeographic inferred events between ecological clusters were dominated by within-cluster events, especially within clusters 1 and 2. Similarly to inferences related to diversity, Ne, and municipality transitions (**Figures 3BC, 5A**), the turn of the century was characterised by an increase in the frequency of transitions within clusters 1 and 2 (**Figure 6D**). Among the between-cluster transitions, those from 2 into 4 were the most frequent, consistent with the highest inferred probability of cluster transitions and likely related to their spatial proximity (**Figure 6A**). We also summarised between-cluster transition events according to whether the cluster mean value of ecological variables increased or decreased with the transition event (**Figure 6E**). Most transitions took place on a positive gradient among the *Meteo* category (to higher temperature and precipitation), and negative gradients for both the *Land* (to lower agriculture and road density) and *Host* (to lower host densities) categories (**Figure 6E**).

## DISCUSSION

This study reports an eco-phylodynamic approach to *M. bovis* within the hotspot risk area of Portugal, representing a multi-host transmission system at the livestock-wildlife interface within a rich ecological landscape. We focused our analyses on a single, newly identified clade related to the Eu3 clonal complex. This is the first study focusing on Eu3 in Iberia, which lacks epidemiological and evolutionary understanding.

Regarding the entire dataset, the SNP-based maximum likelihood tree discriminated four major clades, corresponding to four different clonal complexes of which Eu2 (73%) and Eu3 (25%) were the most representative. This was in agreement with previous studies reporting the prevalence of clonal complexes in Portugal [11, 16, 19]. The SNP distance analysis highlighted local networks, with the geographic distribution of SNP clades suggesting the natural circulation of particular subclades for long periods of time. The observation of several distantly related lineages present in the same region further suggested the occurrence of long-time past introduction events, whose origin could not be determined but is likely to be from regions across the Spanish border. Alternatively, past spatial segregation of TB subpopulations could have led to similar population structures as the ones observed in this study. Disease maintenance was evident in hunting areas since transmission events occurred between different wildlife hosts for long periods of time, with 2008-2016 being the longest period recovered.

We were able to infer *M. bovis* MRCA of the Eu3 clonal complex to have existed around three decades ago, with a substitution rate similar to previous studies [13, 14, 15]. The MRCA was inferred to have been associated with bovine, with subsequent host transition events towards wildlife species, with some reverse host transition from wildlife reservoirs back to bovine also occurring later. There was thus support for both intra- and inter-species transmission routes between all host species, reinforcing the occurrence of historical and sustained cross-species transmission within this ecosystem [5]. The phylogeographic inference also revealed that the MRCA was associated with the Portalegre district (probably from Castelo de Vide municipality) with subsequent geographic transition events towards the Castelo Branco district, and reverse host transition from Castelo Branco back to Portalegre also occurring more recently. Although not possible to demonstrate with the current dataset, this seemingly free geographic transmission range may likely occur due to animal movement (namely wildlife), bovine trading, or both.

Phylogeographic inferences using ecological clusters revealed that transmission events between different ecological settings were dominated by events between two clusters: clusters 2 and 4, located in Portalegre and on the border between Portalegre and Castelo Branco, respectively. There was high support for both symmetric and asymmetric probabilities between the clusters. However, assuming asymmetric probabilities resulted in most events being estimated from cluster 2 into 4. Cluster 2 was the most involved in all inferred cluster transmission events adding to 48% of the total, being also part of 79% of between-cluster events. While our analysis remained purely qualitative without the capacity to infer causation, its outputs were compatible with the notion that wild animals foraging into or away from cluster 2, which is dominated by areas of higher bovine, agriculture, and road densities, could play a role in TB dissemination in this hotspot area. Indeed, between-cluster events were associated with measurable ecological gradients: a positive gradient among climate-related variables towards higher temperature and precipitation, a negative “humanized” gradient (towards higher forest, lower agriculture, lower road density), and a negative gradient of host densities (from high to low density among most hosts). Altogether, these results stress that TB is an established wildlife disease without strict geographic or ecological barriers.

Given that bovine populations are far more restricted in movement than wildlife populations, and in light of the inferred transmission events sourced at ecological cluster 2 in Portalegre towards cluster 4 bordering and including Castelo Branco, these results suggest that existing TB control measures are insufficient to prevent long-distance transmission and spillover within the livestock-wildlife interface. These observations unravel the bordering region between the two districts, within the much larger officially declared hotspot risk area, as a potential corridor of unrecognised importance for the maintenance of *M. bovis*. Indeed, they support the importance of wildlife species in *M. bovis* dissemination in the Portuguese hotspot risk area and the need for new interventions targeting livestock-wildlife interactions. For example, the population expansion of both wild boar and red deer in Europe, including in the Iberian Peninsula [20, 21], is a known risk factor for disease expansion due to animal dispersion, higher animal densities, and aggregation, together with the sympatric character of both species. New strategies focusing on these wildlife reservoirs should be considered to substantially improve the control of animal TB in Portugal. In this area, investment in upscaling sampling and WGS efforts will be critical, as it is the main way to improve data-driven reconstruction of local spatio-temporal histories of *M. bovi*s, essential to design such control strategies.

In sum, the Iberian Peninsula as a whole requires new, improved, and innovative methods to inform science-based management decisions in the ecological interfaces between livestock and wildlife to prevent the overflow of TB and other epidemics.

So far, the implemented measures appear insufficient to prevent between-species transmission across the wildlife-livestock interface. Thus, an improvement of management actions towards reducing contacts and mixing in this interface is critical if aiming to achieve prevalence levels that may sustain strict control or even eradication. This is particularly relevant at first instance for livestock, as it remains the most accessible and manageable host population within which additional control measures may lead towards eradication. Together, increased surveillance of livestock areas adjacent to wildlife hotspot areas or highly dense areas needs to occur due to the detected high transition rates within cattle. Moreover, clear measures to reduce and monitor both livestock and wildlife movements, which seem to be highly responsible for transmission, need to be rigorously taken, particularly in extensive livestock-producing areas and managed hunting areas. Finally, a strong investment effort is needed to find answers to critical questions on which effective control measures depend upon, namely further studies need to be conducted to clarify: the role of other species as intermediate hosts; animal movements and contact patterns as important epidemiological links; and ecological wildlife parameters, such as animal density; and environmental parameters, such as bioclimatic variables, that may influence transmission and TB spread. Our study contributes to this fragmented landscape of control and surveillance by identifying for the first time the bordering region between Portalegre and Castelo Branco as playing a central role in local and long-range dissemination, and perhaps extra priority efforts should first be focused on this region within the much larger official hotspot risk area of Portugal.

## METHODS

### Sampling

*M. bovis* isolates (n=170) were sampled from bovine (n=51), red deer (n=66), and wild boar (n=53) from the hotspot risk area of mainland Portugal (districts: Castelo Branco [n=117], Portalegre [n=53]) bordering Spain (**Supplementary Table T1S1**). Sampling was performed under an official context according to national legislation (see [16] for details). Thirty-six isolates had been previously published [16], with sequence reads obtained from the National Centre for Biotechnology Information SRA database, deposited under accession number PRJNA682618 (accessed: 1 June 2021). The remaining sequences were generated *de novo*.

### DNA Extraction

Novel *M. bovis* isolates (n=134) were stored at -80°C in the National Reference Laboratory of Animal Tuberculosis (INIAV, IP), re-cultured and genomic DNA extracted as in [16].

### Sequence Curation

Genomic DNA was commercially sequenced (Eurofins, Konstanz, Germany) using the Illumina NovaSeq platform (paired-end 150 bp), according to the manufacturer’s specifications. Raw read FASTQ files of all 170 isolates were used for quality control evaluation, trimmed, filtered, and taxonomically classified as in [22]. Samples were analyzed using the vSNP pipeline available at https://github.com/USDA-VS/vSNP (accessed: 1 June 2021) as in [16]. The average depth and genome coverage were 272.7x and 99.69% (respectively). The SNP alignment had a total of 4092 polymorphic positions (**Supplementary Table T1S2**).

### Transmission Mapping

The SeqTrack R library [23, 24] was used to infer local transmission networks through genomic distance minimization between isolates and to keep sampling dates coherent. Transmission trees were constructed with cases grouped within the same transmission chain using a cut-off value of five SNPs within a time frame of five years [25].

### Phylogenomic Analyses

A maximum likelihood method defined with GTR gamma-distributed with invariant sites and four discrete gamma categories, including 1000 bootstrap inferences, was implemented in MEGA-X v10.1.8 [26]. Lineage identification was carried out using KvarQ version 0.12.2 [27] for the assessment of lineage-specific SNPs as in [18].

Temporal signals were measured with TempEst v1.5.3 [28] and LSD v0.3-beta [29]. The best-fitting nucleotide substitution model was selected by Bayesian information criteria (BIC) implemented in jModeltest2 v2.1.10 [30]. Several Bayesian coalescent MCMC analyses were performed in BEAST2 v2.6.2 [31] using the best-fitting nucleotide substitution model together with three molecular clock models and three coalescent demographic priors, resulting in nine different models.

A DATM analysis [32] was implemented in BEAST2 using host species as a discrete trait (bovine, red deer, and wild boar), for both symmetric and asymmetric analysis using the best fitting model (TIM2 with relaxed exponential clock and Bayesian skyline population).

Full details in **Supplementary Text 1**.

### Ecological Clustering

With reference to sample geographical location, twelve ecological variables were sourced from publicly available databases and associated with each sample. We used a heuristic approach based on classic ecology methods (Ward’s clustering, fusion levels) to define groups of samples sharing similar background ecological environments (here termed ecological clusters). To intuitively explore the relationship of ecological clusters with identified lineages and transmission events, we performed multiple factor analysis (MFA, based on principal component analysis) by grouping variables into three categories: *host, land*, and *meteo* (i.e. related to host-species, landscape, and climate, respectively).

Full details on variables used and a step-by-step description of the heuristic clustering and MFA are in **Supplementary Text 2**.

### Phylogeographic Analyses

We performed both geographic and ecological phylodynamic analyses. For geographic analyses, we separately used discrete (sample district or municipality) and continuous (sample latitude-longitude) variables. For ecological analyses, we used the assigned sample cluster as a discrete variable (see Ecological Clustering). Phylogeographic inferences used a coalescent DATM with both symmetric and asymmetric assumptions with the best fitting model (see Phylogenomic Analyses).

Full details in **Supplementary Text 1**.

## SUPPLEMENTARY INFORMATION

We provide supplementary files with supporting material, including Supplementary Text 1 and Text 2 in PDF format (file Supplementary Text.pdf), Supplementary Tables T1S1, T1S3, T1S4 in Excel format (files TableT1S1.xlsx, TableT1S3.xlsx, TableT1S4.xlsx, respectively).

## DATA AVAILABILITY

The sequence data included in this work are deposited under Bioproject accession numbers PRJNA682618 and PRJNA946560 at a public domain server in the National Centre for Biotechnology Information (NCBI) SRA database. All the tables describing the samples and their metadata are provided as supplementary Excel files.

## ACKNOWLEDGEMENTS

MVC acknowledges funding from Fundação para a Ciência e a Tecnologia, IP (FCT)/MCTES through national funds (PIDDAC) and co-funding by the European Regional Development Fund (FEDER) of the European Union, through the Lisbon Regional Operational Program and the Competitiveness and Internationalisation Operational Program for Portugal 2020 or other programs that may succeed in the scope of the project “Colossus: Control Of tubercuLOsiS at the wildlife/livestock interface uSing innovative natUre-based Solutions” (references PTDC/CVT-CVT/29783/2017, LISBOA-01-0145-FEDER-029783, POCI-01-0145-FEDER-029783). Strategic funding from FCT to cE3c and BioISI Research Units (UIDB/00329/2020 and UIDB/04046/2020) and to the associate laboratory CHANGE (LA/P/0121/2020) are gratefully acknowledged. ACP was funded by FCT (SFRH/BD/136557/2018).

## Supplementary information captions

**Supplementary Text T1** – Sequencing and phylogenetics.

**Table T1S1** - Metadata of all 170 *M. bovis* isolates under study.

**Table T1S2** – Sequencing statistics of all 170 *M. bovis* isolates used in this study.

**Table T1S3** – Temporal signal analysis by root-to-tip test using TempEst.

**Table T1S4** - Analysis of best-fitting nucleotide substitution model using jModelTest2.

**Table T1S5** – Path sampling analysis for the selection of the best-fitting phylogenomic model in BEAST2 by marginal likelihood estimator and Bayes factor comparison.

**Table T1S6** – Path sampling analysis for the selection of the best-fitting ancestral trait analysis model, for both host trait and phylogeographic analysis in BEAST2 by marginal likelihood estimator and Bayes factor comparison.

**Figure T1S1** – *In silico* spoligotype patterns of *M. bovis* population in Portugal.

**Figure T1S2** – Temporal signal analysis of European 3 clonal complex.

**Figure T1S3** – Phylogeographic analysis of European 3 clonal complex using district as a grouping factor.

**Figure T1S4** – Phylogeographic analysis of European 3 clonal complex.

**Figure T1S5** – Posterior probabilities of geographic transitions of European 3 clonal complex.

**Supplementary Text T2** – Ecological clustering analyses

**Figure T2S1** – Ward’s minimum variance clustering dendrogram.

**Figure T2S2** – Fusion level values for the dendrogram presented in Figure T2S1.

**Figure T2S3** – Ward’s minimum variance clustering dendrogram.

**Figure T2S4** – Number of samples per cluster when considering sets of 4 or 5 clusters.

**Figure T2S5** – Multiple factor analysis output.

**Figure T2S6** – Multiple factor analysis variable contribution to dimensions 1 and 2.

**Figure T2S7** – Sample clusters and MFA top variable visualization.

**Figure T2S8** – Sample clusters in MFA space (top contributing variables) colored by variable variation.

