## Supplemental Text and Figures for "Population structure and history of *Mycobacterium bovis* European 3 clonal complex reveal transmission across ecological corridors of unrecognised importance in Portugal"

This supplementary material is hosted by *Emerging Microbes and Infection* as supporting information alongside the article “**Population structure and history of *Mycobacterium bovis* European 3 clonal complex reveal transmission across ecological corridors of unrecognised importance in Portugal”**. The same standards for ethics, copyright, attributions and permissions as for the article apply.

### Supplementary Text T1 – Sequencing and phylogenetics (pages 3-10)

**Additional methods description**

**Additional results**

**Supplementary references**

**Table T1S3** – Temporal signal analysis by root-to-tip test using TempEst.

**Table T1S5** – Path sampling analysis for the selection of the best-fitting phylogenomic model in BEAST2 by marginal likelihood estimator and Bayes factor comparison.

**Table T1S6** – Path sampling analysis for the selection of the best-fitting ancestral trait analysis model, for both host trait and phylogeographic analysis in BEAST2 by marginal likelihood estimator and Bayes factor comparison.

**Figure T1S1** – *In silico* spoligotype patterns of *M. bovis* population in Portugal.

**Figure T1S2** – Temporal signal analysis of European 3 clonal complex.

**Figure T1S3** – Phylogeographic analysis of European 3 clonal complex using district as a grouping factor.

**Figure T1S4** – Phylogeographic analysis of European 3 clonal complex.

**Figure T1S5** – Posterior probabilities of geographic transitions of European 3 clonal complex.

### Supplementary Text T2 – Ecological clustering analyses (pages 11-18)

**Data variables**

**Data sources**

**Heuristic approach for selection of N possible ecological clusters**

**Figure T2S1** – Ward’s minimum variance clustering dendrogram.

**Figure T2S2** – Fusion level values for the dendrogram presented in Figure T2S1.

**Figure T2S3** – Ward’s minimum variance clustering dendrogram.

**Figure T2S4** - Number of samples per cluster when considering sets of 4 or 5 clusters.

**Phylogeography**

**Multiple factor analysis of N=4 ecological clusters**

**Figure T2S5** - Multiple factor analysis (MFA) output.

**Figure T2S6** - MFA variable contribution to dimensions 1 and 2.

**Figure T2S7** - Sample clusters and MFA top variable visualization.

**Figure T2S8** - Sample clusters in MFA space (top contributing variables) colored by variable variation.

**Supplementary references**

### Supplementary Text T1 – Sequencing and phylogenetics

#### Additional methods description

**Phylodynamic analyses**

The temporal signal of the dataset’s phylogenetic tree, the identified clades, and isolates grouped by host species and geographic location was explored with TEMPoral Exploration of Sequences and Trees software (TemEst version 1.5.3) (1) by a root-to-tip test, evaluated via the positive correlation (r^2^) between SNP distance and time. One representative clade was selected for further phylogenomic and phylogeographic inferences based on its stronger temporal signal: clade 10 (all Eu3 clonal complex isolates; n=44). For this clade, the Least-Squares Dating software (LSD, version 0.3-beta) (2) was used to evaluate the temporal signal more robustly by a date-randomization test (3).

Several Bayesian coalescent Markov Chain Monte Carlo (MCMC) analyses were performed in BEAST2 version 2.6.2 (4) using the best-fitting nucleotide substitution model together with three molecular clock models (strict, relaxed exponential, and relaxed log normal), and three coalescent demographic priors (constant, exponential, and Bayesian skyline population), resulting in nine different models (5-8). Independent MCMC analyses were run for 50 million generations steps and posterior distributions were sampled every 5000 generations. For all models, two runs were performed, and model parameters were assessed for convergence and satisfactory effective sample sizes (ESS >200) in Tracer version 1.7.1 (9). The runs were combined in LogCombiner version 2.6.6 (10), where trees were also subsampled, and a maximum clade credibility tree was created with median heights (10% burn-in) using TreeAnnotator version 2.6.3 (10) and visualised in FigTree version 1.4.4 (10). Model performance was evaluated by maximum likelihood estimation (MLE) based on path sampling (PS) (11) and paired comparisons (of all models to the first combination: TIM2, strict clock, and constant population) of marginal likelihoods using Bayes Factors (BF) (12). This approach was used in all phylodynamic analyses (i.e. original phylogenomic analysis, ancestral state host reconstruction using discrete traits, and phylogeographic analysis). The relative estimated posterior probabilities (PP) of transition between host species, districts, municipalities, and ecological clusters, from both symmetric and asymmetric models, both also in the MASCOT and MultiTypeTree models, were obtained.

Lineages-through-time analysis and coalescent Bayesian Skyline analysis were performed in Tracer using the best-fitting model to measure strain differentiation and assess population dynamics during the study period.

**Phylogeography analyses**

A polyphasic approach was taken addressing phylogeographic inferences, either using discrete (districts, municipalities) or continuous (latitude, longitude coordinates) geographic data. For discrete data, three types of models were used: coalescent MCMC (i.e. DATM analysis using both symmetric and asymmetric assumptions), structured coalescent MCMC by a MultiTypeTree model (13), and approximate structured coalescent MCMC by a MASCOT model (14). In phylogeographic analyses, continuous MCMC models, as well as the MASCOT and MultiTypeTree models when municipalities were used as discrete traits, did not converge. The backward migration rates of transition between geographic locations using districts as discrete traits were also obtained.

Spatial phylogenetic reconstruction of evolutionary dynamics (SPREAD version 1.0.7) (15) was used to summarise and compute discrete BF for DATM models, with the ones with BF >3 being considered of high occurrence probability. Transitions between municipalities inferred by the phylogeographic coalescent asymmetric model, as well as transitions between ecological clusters inferred by the phylogeographic coalescent model were recovered from internal nodes.

#### Additional results

**Phylogenetic analysis**

*In silico* spoligotyping was performed to assess genetic diversity and to compare with wet-lab spoligotyping results (**Supplementary Table T1S2**). We identified 19 different spoligotypes, with SB1174 (25%), SB0122 (19%), SB0121 (14%), SB1264 (12%), and SB1195 (8%) being the top five profiles and accounting for 79% of total isolates (**Supplementary Figure T1S1A**). These five spoligotypes were registered in all three host species (bovine, red deer, wild boar), with prevalence rates varying across geographic regions and host species. The spoligotype SB0122 (26%) was commonly found in Castelo Branco, while SB1174 (49%) was dominant in Portalegre (**Supplementary Figure T1S1B**). SB1174 was the most reported in bovine (20%) and wild boar (22%), while SB0122 (24%) was dominant in red deer (**Supplementary Figure T1S1C**). A total of 85% of spoligotypes were concordant with wet-lab spoligotypes, with the majority of discordant cases (79%) resulting from changes in one or two spacers.

For 24 isolates, the spoligotyping profile obtained by the reverse-hybridization method was discordant with the profile determined *in silico*, a finding reported by others (16, 17). Considering the top 5 spoligotypes, no host or geographic-specific spoligotype was determined, supporting inter-host species and inter-district transmission patterns.

**Temporal signal evaluation**

Before the Bayesian phylogenomic analysis, the temporal signal of the *M. bovis* population was assessed. We used a TempEst by root-to-tip analysis, which tests the correlation between sampling time and genomic divergence (SNP differences), performed by considering the entire population, dividing it by phylogenetic clades, geographic regions, and host species (Supplementary Table T1S3). Based on the entire population, no temporal signal was found (r^2^=0.005) and only a few subpopulations showed a positive correlation (r^2^>0.1), namely, phylogenetic clades 1, 6, 7, and 10.

**Phylogeography analyses**

To explore the geographic relationship between the Eu3 clonal complex isolates, phylogeographic inferences were performed to estimate the internal node probability relating to geographic location. Two main phylogeographic inferences were attempted: the first using discrete traits (district, municipality) and the second using continuous traits (latitude, longitude). Only the first attempt was successful since continuous trait models did not converge. Three types of models were tested: coalescent, structure coalescent, and approximated structure coalescent models. Both structure and approximated structure coalescent models did not converge, likely due to the high number of municipalities (N=8).

The inferred transition probability between both districts was close to 1.00 (**Supplementary Figure T1S3**), meaning that the entire area of study could be interpreted as a geographic unit. Considering the less supported asymmetric coalescent model (**Supplementary Figure T1S4**), high probabilities of transition between districts in both directions were also estimated (Castelo Branco – Portalegre, PP=0.97, BF=1630.91; Portalegre – Castelo Branco, PP=1.00, BF=7.00) (**Supplementary Figure T1S3**), supporting the study area as a single geographic unit hypothesis.

Both the structure and approximated structure coalescent models have the additional assumption, compared to the simple coalescent model, that the transition between states is more unlikely than the transmission within states, forcing a structure within the population (**Supplementary Figure T1S4**). Worth noting that approximated structure coalescent models (using MASCOT) showed relatively low migration rates (0.05; **Supplementary Figure T1S3**), but the structure coalescent model (by MultiTypeTree) showed a relatively high migration rate from Portalegre to Castelo Branco (0.83) with an almost nonexistent migration rate in the other direction (7.5 x 10^-12^) (**Supplementary Figure T1S3**).

Furthermore, coalescent models using municipalities as grouping levels showed relatively low transition probabilities, in both symmetric and asymmetric models, with only five transitions showing relatively high probabilities: in the symmetric model, only within districts transition showed high probabilities (Castelo Branco – Idanha-a-Nova, PP=1.00, BF=4.24; Castelo de Vide – Nisa, PP=0.63, BF <3.00; Castelo de Vide – Portalegre, PP=0.62, BF <3.00; Nisa – Portalegre, PP=0.64, BF <3.00) (**Supplementary Figure T1S5**); in the asymmetric model, only one transition showed high probability, being between districts (Idanha-a-Nova – Avis, PP=0.90, BF=3.78) (**Supplementary Figure T1S5**).

**Supplementary Table T1S3** – Temporal signal analysis by root-to-tip test using TempEst.

| Group | Total | Cattle | Wild boar | Red deer | Wildlife | Castelo Branco | Portalegre | 1 | 2 | 5 | 6 | 7 | 10 |
| --- | --- | --- | --- | --- | --- | --- | --- | --- | --- | --- | --- | --- | --- |
| Date range | 16 | 16 | 12 | 14 | 14 | 16 | 15 | 16 | 10 | 11 | 12 | 15 | 15 |
| Slope (rate) | 0.0009 | 0.7437 | 0.1834 | 0.0024 | 0.1338 | 0.1234 | 0.0030 | 0.0001 | 0.0001 | 0.0006 | 0.0002 | 0.0011 | 0.0003 |
| X-Intercept (TMRCA) | 1917 | 1982 | 1983 | -200 | 2001 | 2002 | 870 | 1990 | 1908 | 1974 | 1994 | 1980 | 1991 |
| R squared | **0.005** | **0.066** | **0.003** | **0.061** | **0.023** | **0.022** | **0.028** | **0.253** | **0.007** | **0.032** | **0.303** | **0.236** | **0.200** |

**Supplementary Table T1S5** – Path sampling analysis for the selection of the best-fitting phylogenomic model in BEAST2 by marginal likelihood estimator and Bayes factor comparison.

| Nucleotide Substitution Model | Molecular Clock Model | Coalescent Demographic Priors | Marginal Likelihood Estimator | Bayes Factor |
| --- | --- | --- | --- | --- |
| TIM2 + I + G | Strict | Constant Population | -11828.93 | - |
| TIM2 + I + G | Strict | Exponential Population | -9961.10 | 1867.83 |
| TIM2 + I + G | Strict | Bayesian Skyline | -11080.19 | 748.74 |
| TIM2 + I + G | Relaxed Exponential | Constant Population | -10080.56 | 1748.37 |
| TIM2 + I + G | Relaxed Exponential | Exponential Population | -10056.95 | 1771.98 |
| TIM2 + I + G | **Relaxed Exponential** | **Bayesian Skyline** | **-9820.09** | **2008.84** |
| TIM2 + I + G | Relaxed Log Normal | Constant Population | -10126.61 | 1702.32 |
| TIM2 + I + G | Relaxed Log Normal | Exponential Population | -10099.97 | 1728.96 |
| TIM2 + I + G | Relaxed Log Normal | Bayesian Skyline | -10005.31 | 1823.62 |

**Supplementary Table T1S6** – Path sampling analysis for the selection of the best-fitting ancestral trait analysis model, for both host trait and phylogeographic analysis in BEAST2 by marginal likelihood estimator and Bayes factor comparison.

| Model Trait | Model Type | Marginal Likelihood Estimator | Bayes Factor |
| --- | --- | --- | --- |
| Host | **Symmetric** | **-10116.59** | **-** |
| Host | Asymmetric | -10150.55 | -33.96 |
| District | **Symmetric** | **-10105.60** | **-** |
| District | Asymmetric | -10139.37 | -33.77 |
| District | MASCOT | -10361.37 | -255.77 |
| District | MultiTypeTree | -10578.21 | -472.61 |
| Municipality | Symmetric | -10218.75 | -113.15 |
| Municipality | **Asymmetric** | **-10162.21** | **-56.61** |
| Ecological Cluster 4 | **Symmetric** | **-10099.99** | **5.61** |
| Ecological Cluster 4 | **Asymmetric** | **-10122.23** | **-16.63** |
| Ecological Cluster 5 | Symmetric | -10178.79 | -73.19 |
| Ecological Cluster 5 | Asymmetric | -10273.86 | -168.26 |

**Supplementary Figure T1S1 –** *In silico* spoligotype patterns of *M. bovis* population in Portugal. The total number of each spoligotype pattern (A) is represented, together with the top 5 spoligotype patterns divided by geographic region (B) and by host species (C).

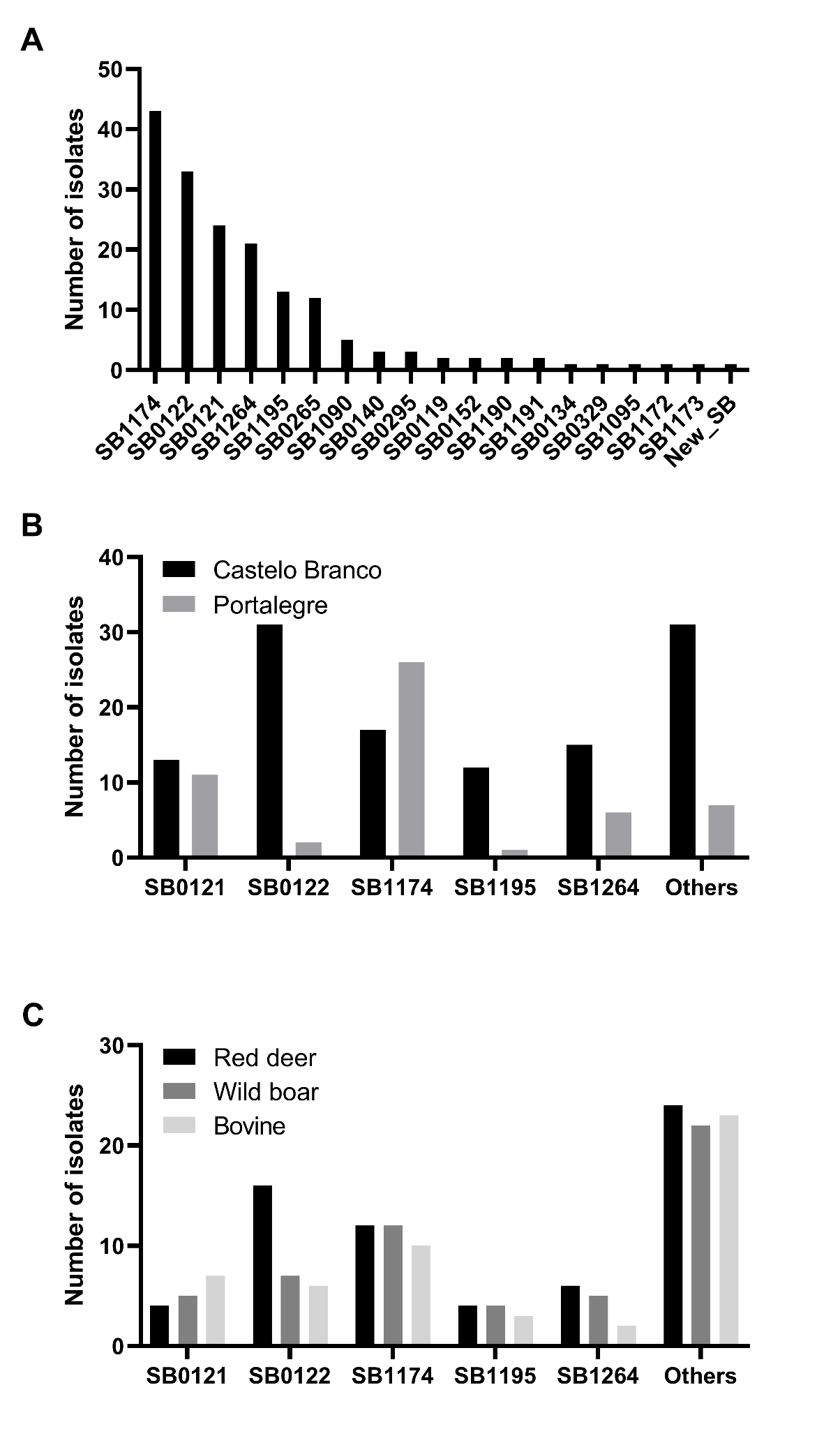

**Supplementary Figure T1S2** – Temporal signal analysis of European 3 clonal complex. Least-Squares Dating software was used to perform a date-randomization test.

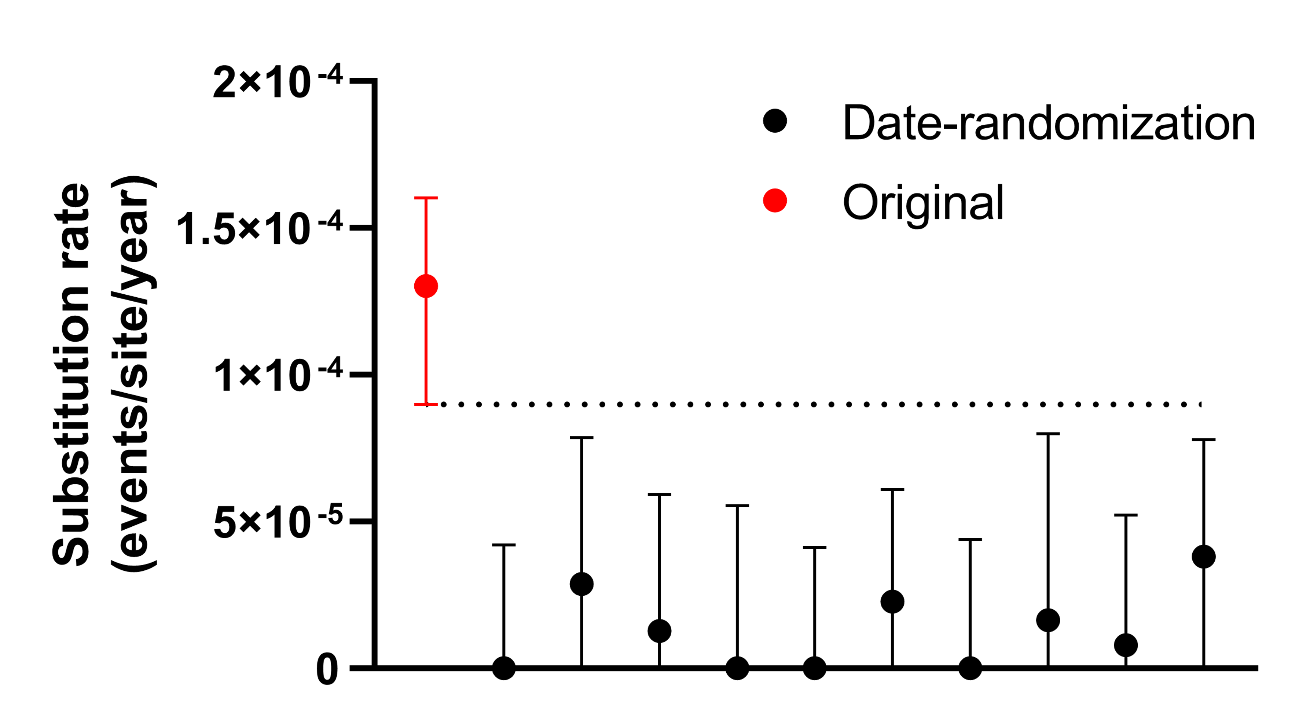

**Supplementary Figure T1S3** – Phylogeographic analysis of European 3 clonal complex using district as a grouping factor. A) Maximum credibility tree was estimated under a model of symmetric district transitions with posterior support for major nodes shown with grey bars indicating the 95% highest posterior density intervals for node date estimates. Districts are colour labels (Castelo Branco = red, Portalegre = blue). B) District state posterior probabilities under a coalescent model of symmetric and asymmetric transitions. C) District state backward migration rates under an approximated structure coalescent model (MASCOT) and structure coalescent model (MultiTypeTree).

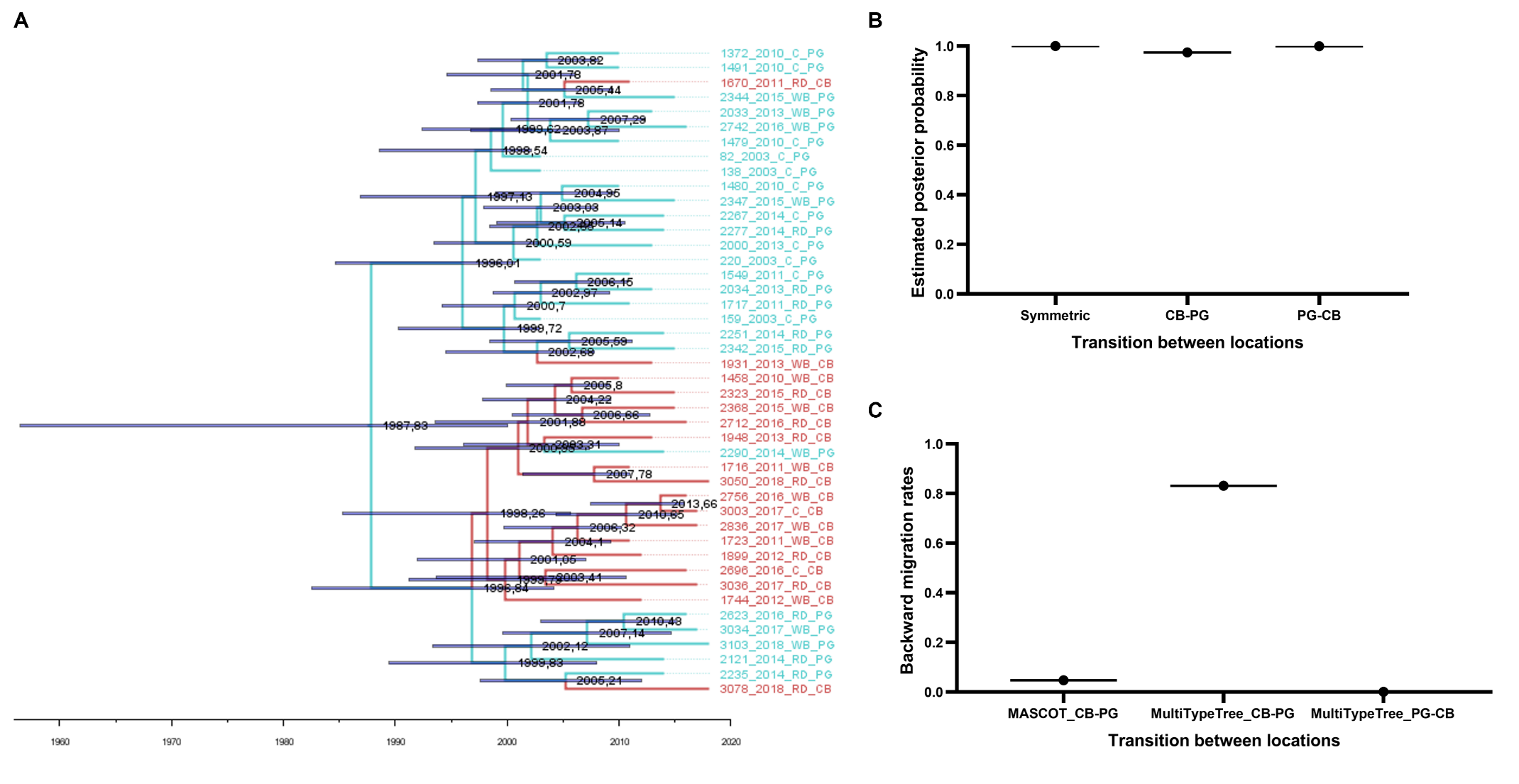

**Supplementary Figure T1S4** – Phylogeographic analysis of European 3 clonal complex. Using district as a grouping factor, three models were performed: asymmetric (A), approximated structure (B), and structure (C). Using municipality as a grouping factor, an asymmetric model was also performed (D).

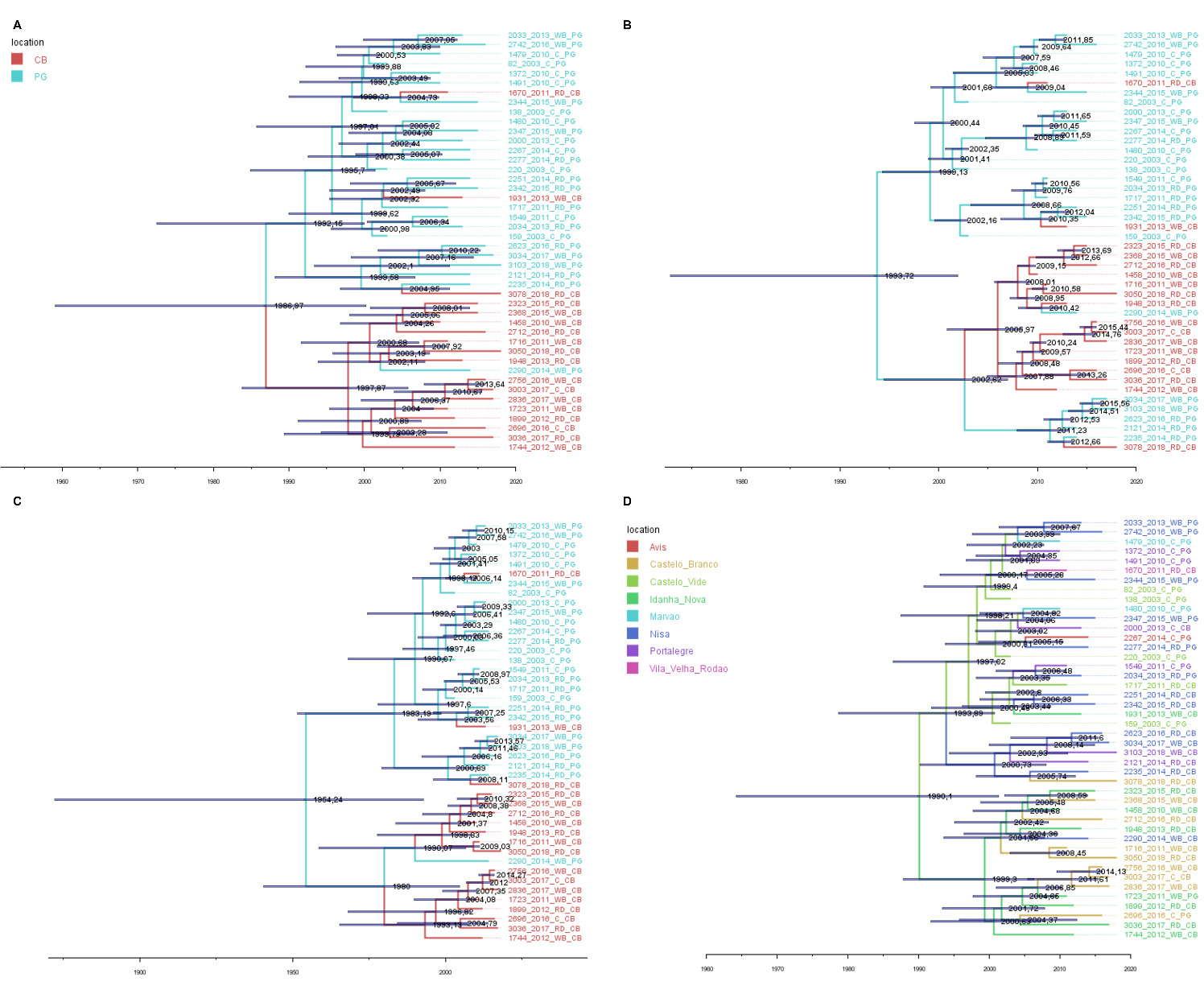

**Supplementary Figure T1S5 –** Posterior probabilities of geographic transitions of European 3 clonal complex. Using municipality as a grouping factor, both estimated posterior probabilities for symmetric (A) and asymmetric (B) models were performed. Colors highlight posterior probabilities >0.5.

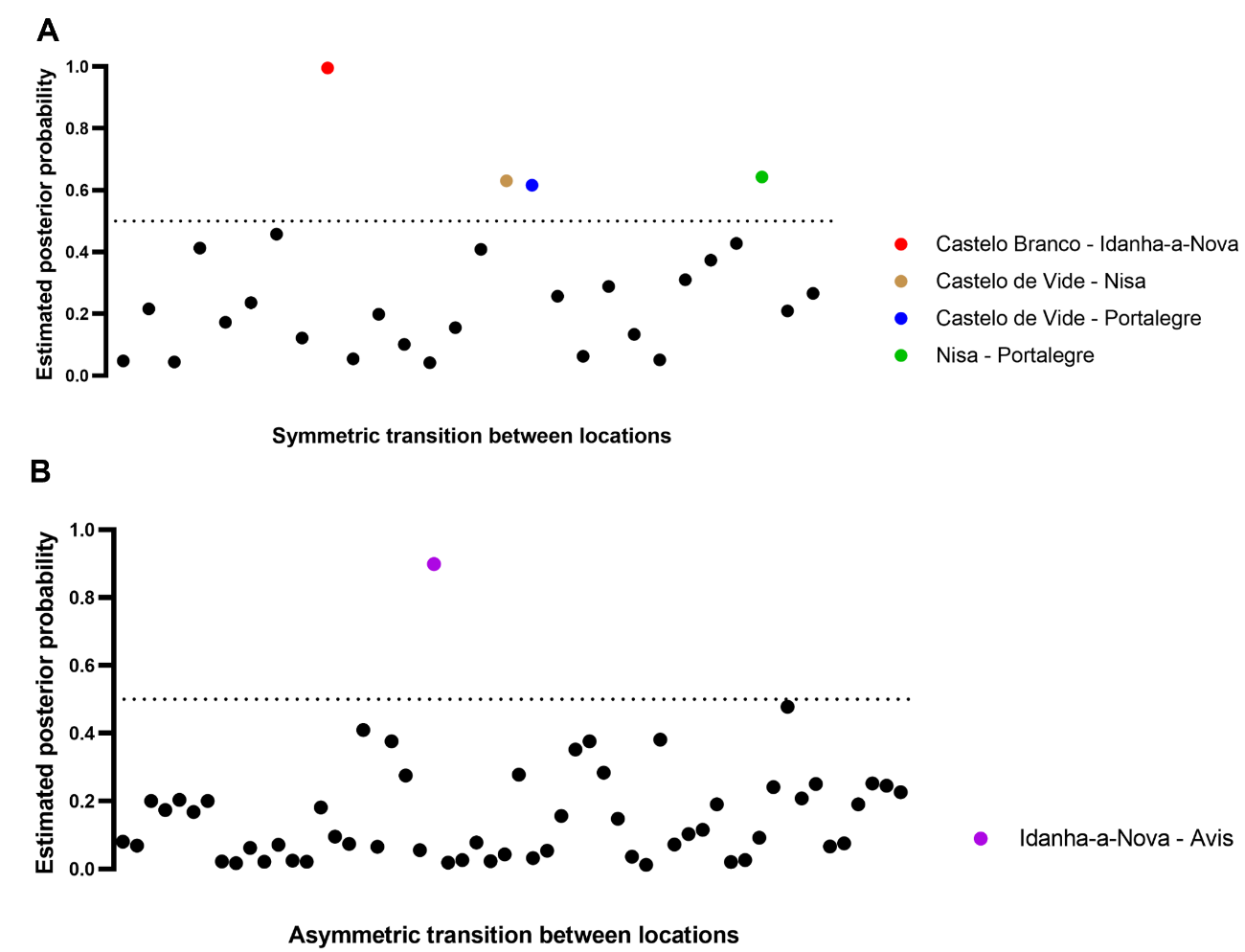

### Supplementary Text T2 – Ecological clustering analyses

#### Data variables

**Table T2S1** – Summary of environmental variables used.

| **density_human** | number of humans per municipality / normalized by area |
| --- | --- |
| **density_bovine** | number of bovine per municipality / normalized by area |
| **density_ovi** | number of animal (Mufflon) hunted per municipality / normalized by area |
| **density_dam** | number of animal (Fallow deer) hunted per municipality / normalized by municipality area |
| **density_sus** | number of animal (Wild boar) hunted per municipality / normalized by municipality area |
| **density_cer** | number of animal (Red deer) hunted per municipality / normalized by municipality area |
| **agriculture** | Agriculture coverage (%) |
| **forest** | Forest coverage (%) |
| **roads_t** | Road network density |
| **temp_ann** | Average annual temperature (ºC) |
| **prec_ann** | Average annual precipitation (mm) |
| **prec_dry_m** | Driest month precipitation (mm) |

Each variable was averaged across a circular area around the location of each sample within a 2 km radius [[1]](https://paperpile.com/c/moWVJ6/P4ou), in order to cover the widest vital area of the host species considered in this study. When variables were available with a value for each municipality, we calculated a weighted average, using the proportion occupied in the 2 km radius buffer as the weight.

#### Data sources

Land cover data (for variables **agriculture**, **forest**) was retrieved from the Copernicus.eu database [[2]](https://paperpile.com/c/moWVJ6/dhBi). We aggregated available Corine Land Cover (CLC) classes into two categories: agriculture (CLC 200 to 299) and forest (CLC 300 to 399). We used the average extent of cover for both categories considering the years 2000, 2006, 2012 and 2018 (to cover the time range of our samples).

National livestock numbers were retrieved from Instituto Nacional de Estatística (INE, Portuguese National Institute of Statistics) [[3]](https://paperpile.com/c/moWVJ6/8uCa) for the years 2009 and 2019. Since data is officially reported per municipality, we calculated livestock average density per municipality (livestock numbers divided by municipality area). For each sample, using the respective municipality as reference, we set the variable **density_bovine** as the average bovine density per municipality between 2009 and 2019.

Wild ungulates abundance was inferred from hunting bag numbers retrieved from Instituto da Conservação da Natureza e das Florestas (ICNF, Portuguese Institute of Nature and Forests) [personal communication to Mónica V. Cunha]. Data included red deer (*Cervus elaphus*), Fallow deer (*Dama dama*), European mouflon (*Ovis aries*), and wild boar (*Sus scrofa*) numbers for hunting seasons starting 1992/1993 and ending 2017/2018 per municipality. We calculated the hunting density per municipality by dividing bag numbers by municipality area across the years and used this density as a proxy for local abundance. For each sample, using municipality as reference, we set the variable **density_ovi** as the local density abundance of European mouflon, **density_dam** as the density of Fallow deer, **density_sus** as the density of wild boar, and **density_cer** as the density of red deer.

Climate data (for variables **temp_ann**, **prec_ann**, **prec_dry_m**) was obtained from the WorldClim dataset [[4,5]](https://paperpile.com/c/moWVJ6/8KOf+X8Hv). Monthly historical climate data was downloaded from the WorldClim website for the years 2000, 2006, 2012 and 2018, in order to match the years for which Land Cover data was available (see details above). Using the biovars function in the ‘dismo’ package in R (v1.3-8) [[6]](https://paperpile.com/c/moWVJ6/pilw), we calculated the average for the variables **temp_ann**, **prec_ann** and **prec_dry_m** for each sample.

Human infrastructure data (for variable **roads_t**) was obtained from the Global Roads Inventory Project (GRIP) dataset [[7]](https://paperpile.com/c/moWVJ6/NxyF). Road data density was classified according to five categories: "highways", "primary roads", "secondary roads", "tertiary roads" and "local roads". Data for Portugal and Spain included in GRIP was gathered primarily from OpenStreetMap [[8]](https://paperpile.com/c/moWVJ6/rrf0), and the sum of the categories was used to define **roads_t** per sample.

#### Heuristic approach for selection of N possible ecological clusters

We started our approach by performing variance clustering of the 13 environmental data variables over the 170 samples using the Ward’s minimum variance clustering approach. The objective was to define groups of samples sharing similar environments (from now on termed ecological clusters, or simply clusters) within which the sum of squares is minimized. We used the base function *hclust()* in R, which implements the classic Ward’s method [[9–11]](https://paperpile.com/c/moWVJ6/xibt+T467+iUle). The resulting, clustering dendrogram is presented in **Figure T2S1**.

With the obtained clustering, we proceeded to evaluate at which height should the dendrogram be cut in order to obtain two alternative sets of clusters (i.e. not with too few numbers of clusters, nor with too many clusters). A common heuristic to select an adequate cutting height is to look at Fusion Level Values (FLVs) [[11]](https://paperpile.com/c/moWVJ6/iUle). FLVs represent the dissimilarity levels at which two branches tend to coalesce (fuse) across the dendrogram.

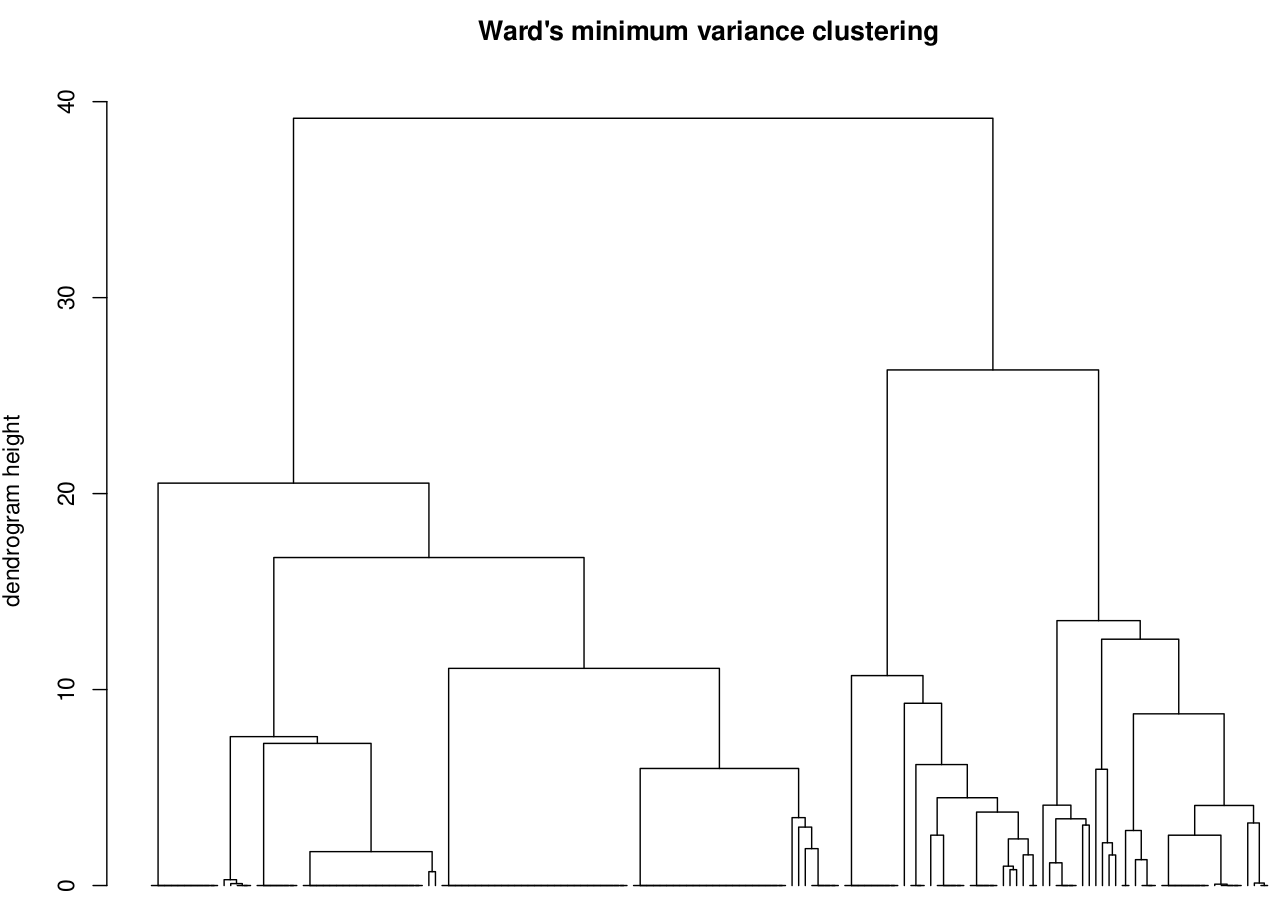

**Figure T2S1 –** Ward’s minimum variance clustering dendrogram.

**Figure T2S2** presents the FLVs for the dendrogram of **Figure T2S1**, showing that as the considered cutting height decreases, the number of resulting clusters increases. For example, a cut between 15 and 20 would always result in 4 clusters. Intuitively, the wider the height range that would always result in the same number of clusters, the “cleaner” the cut is considered to be. For example, to obtain 2 clusters, any height between ~28 and 40 can be considered, which is the widest range in the presented FLVs (widest horizontal segment).

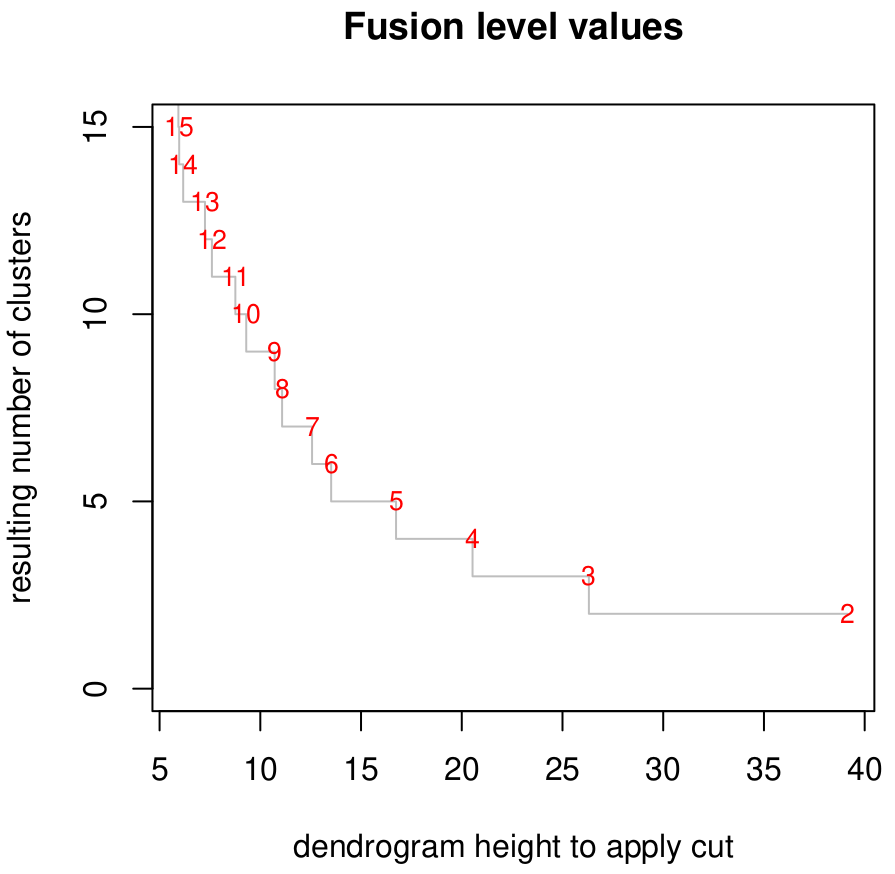

**Figure T2S2 –** Fusion level values for the dendrogram presented in Figure T2S1.

We selected the two cleanest possible cuts that would at the same time result in more than 3 clusters. Cuts with height >21 would result in either 2 or 3 clusters and were thus not considered. The next cleanest cuts were between height 17 and 21 leading to 4 clusters, and between 13 and 17 leading to 5 clusters (**Figures T2S2, T2S3**).

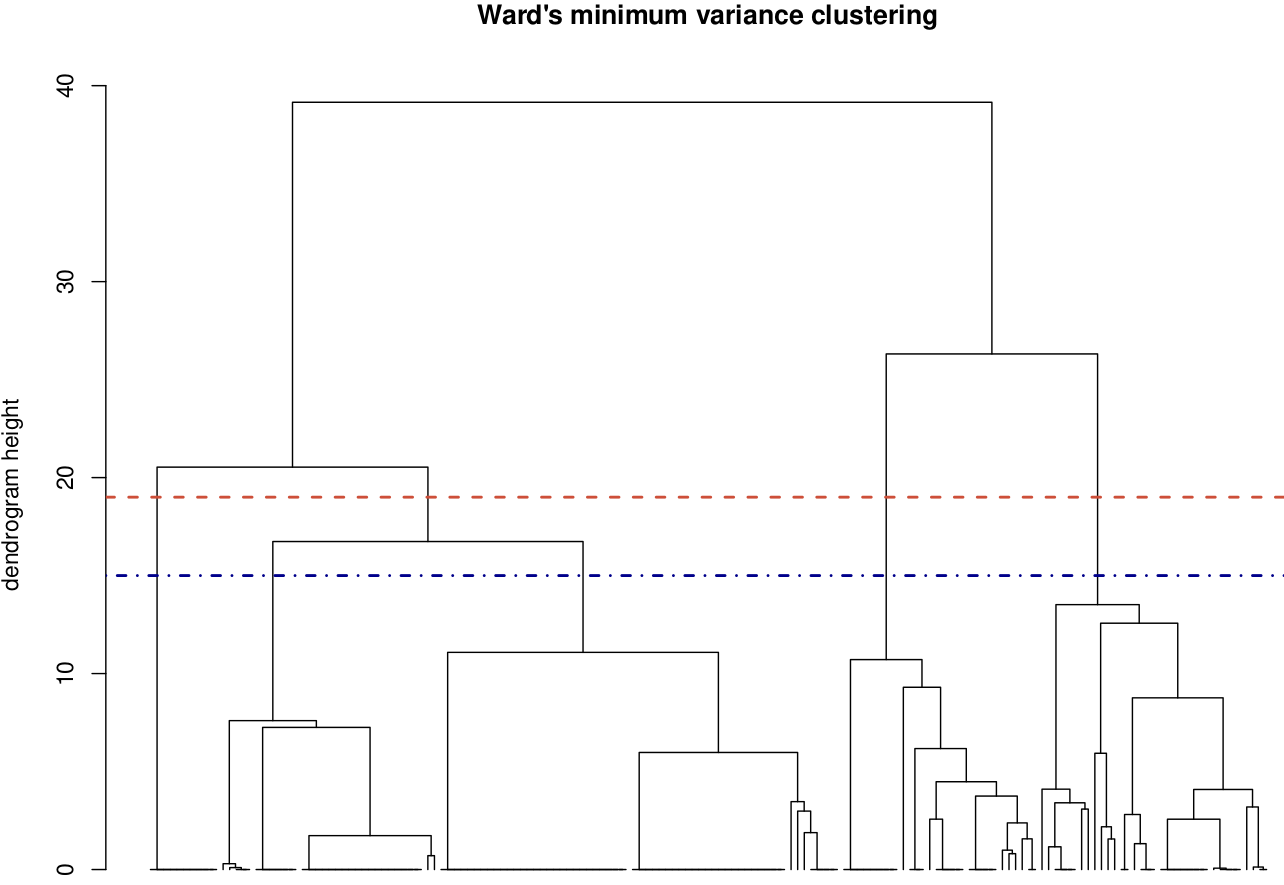

**Figure T2S3 –** Ward’s minimum variance clustering dendrogram. Presented are the height cuts at 15 leading to 5 clusters (dotted-dashed blue horizontal line) and at 19 leading to 4 clusters (dashed red horizontal line).

When considering either the 4 or 5 sets of clusters, the majority of samples belonged to a single cluster (**Figure T2S4**). Apart from this, all other clusters included more than 10 samples (minimum was N=11).

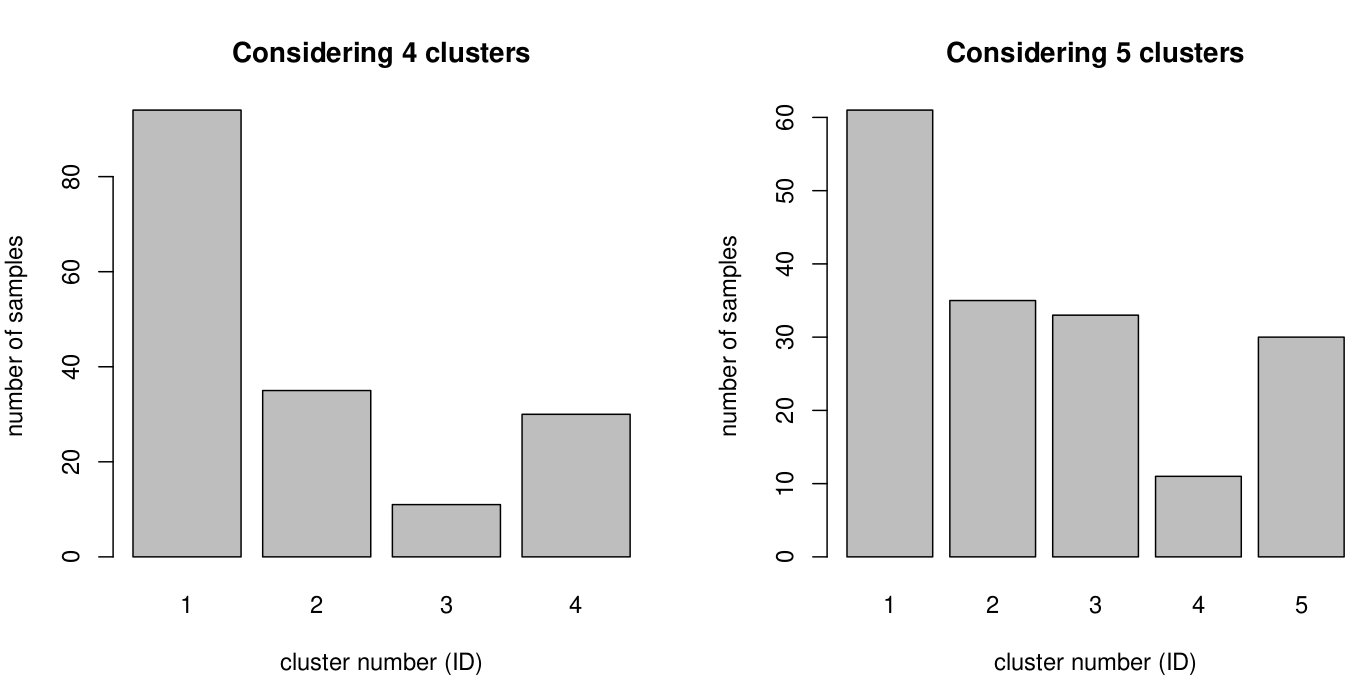

**Figure T2S4** – Number of samples per cluster when considering sets of 4 or 5 clusters.

#### Phylogeography

At this point we tested phylogeography with 4 or 5 clusters and got support for 4 clusters. The rest of the material focuses on applying PCA to the 170 samples and presenting the results in the context of 4 clusters only. Note that the PCA analyses are blind to the number of clusters (it is not an input variable of the PCA method).

#### Multiple factor analysis of N=4 ecological clusters

As described in the phylogeography analyses present in the main text, the clustering of Eu3 clonal complex isolates was tested assuming either 4 or 5 possible geographical clusters. Using 4 clusters (independently of the symmetry assumption used) was the best-fitting model, according to BF analysis. We therefore proceeded with a multiple factor analysis (MFA) using the 4 clusters described above.

MFA is a useful method to explore complex relationships between multiple variables or groups of variables [[11]](https://paperpile.com/c/moWVJ6/iUle). When all variables are quantitative (such as in **Table T2S1**), MFA is in essence a Principal Components Analysis (PCA) applied to the entire set of variables but in which weighting of subsets of variables (groups) can be used. MFA first computes a PCA for each centered group of variables. Each centered table is then weighted, while accounting for different variances among the groups (by dividing by the square root of the first eigenvalue) so that all groups receive equal weights in the following global analysis. The weighted groups of variables are put together and submitted to a global PCA. By comparing the PCA of each group with the global PCA, existing data structures, signatures and differences can be assessed among samples, variables and groups of variables. MFA allows for an exploratory view where correlative structures can be exposed without reference to directionality or causality. We performed MFA using the function *MFA()* from the R-package *FactoMineR* (v2.4) [[12]](https://paperpile.com/c/moWVJ6/cS7O), to which we provide 3 variable groups: Host (***density_human, density_bovine, density_ovi, density_dam, density_sus, density_cer***), Meteo (***temp_ann, prec_ann, prec_dry_m***) and Land (***agriculture, forest, roads***).

**Figure T2S5** illustrates the correlation between variable groups and the two main dimensions of the global PCA. Together, dimension 1 and 2 represented 61.6% of the total variance in the data.

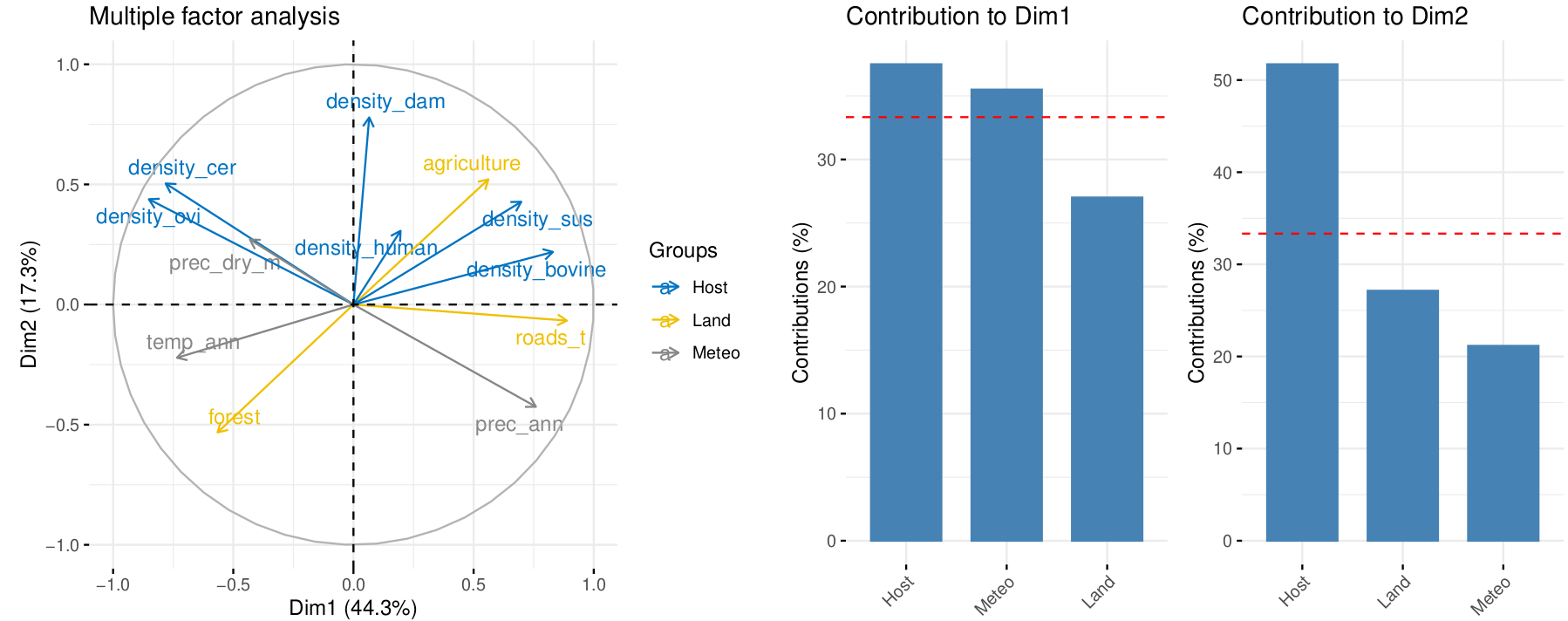

**Figure T2S5 –** Multiple factor analysis (MFA) output. Groups of variables (Host, Land, Meteo) are shown with different colors.

Visual inspection of the global PCA results allowed to gain insights into inherent correlative data structures and signatures (before directly assessing ecological clusters), for example:

- The Host variable group could be interpreted under three subgroups:
  - group 1 (HG1) was characterized by the density of red deer (***density_cer***) and Mufflon (***density_ovi***) grouping together and being positively correlated.
  - group 2 (HG2) was characterized by the density of wild boar (***density_sus***) and Bovine (***density_bovine***) grouping together and being positively correlated.
  - group 3 (HG3) was characterized by the density of fallow deer (***density_dam***) being correlated with the density of human (***density_human***).
- The Meteo variable group could be interpreted under two subgroups:
  - group 1 (MG1) was characterized by precipitation in the driest month (***prec_dry_m***) and annual mean precipitation (***prec_ann***) being negatively correlated.
  - group 2 (MG2) included only annual mean temperature (***temp_ann***).
- The Land variable group could be interpreted under two subgroups:
  - group 1 (LG1) showing that percent of forest (***forest***) and agriculture (***agriculture***) land coverage were negatively correlated.
  - group 2 (LG2) that included only the density of roads (***roads_t***).

As shown in **Figure T2S6**, the largest contributors to dimension 1 (above the uniform expectation) were the mean annual temperature (***temp_ann***), density of roads (***roads_t***), precipitation (***prec_ann***), and densities of Mufflons (***density_ovi***), Bovines (density_bovine) and Red deers (***density_cer***). For dimension 2, the largest contributors (above the uniform expectation) were the density of Fallow deers (***density_dam***), the percent of forest (***forest***) and agriculture (***agriculture***) on land coverage, precipitation (***prec_ann***) and density of Red deers (***density_cer***).

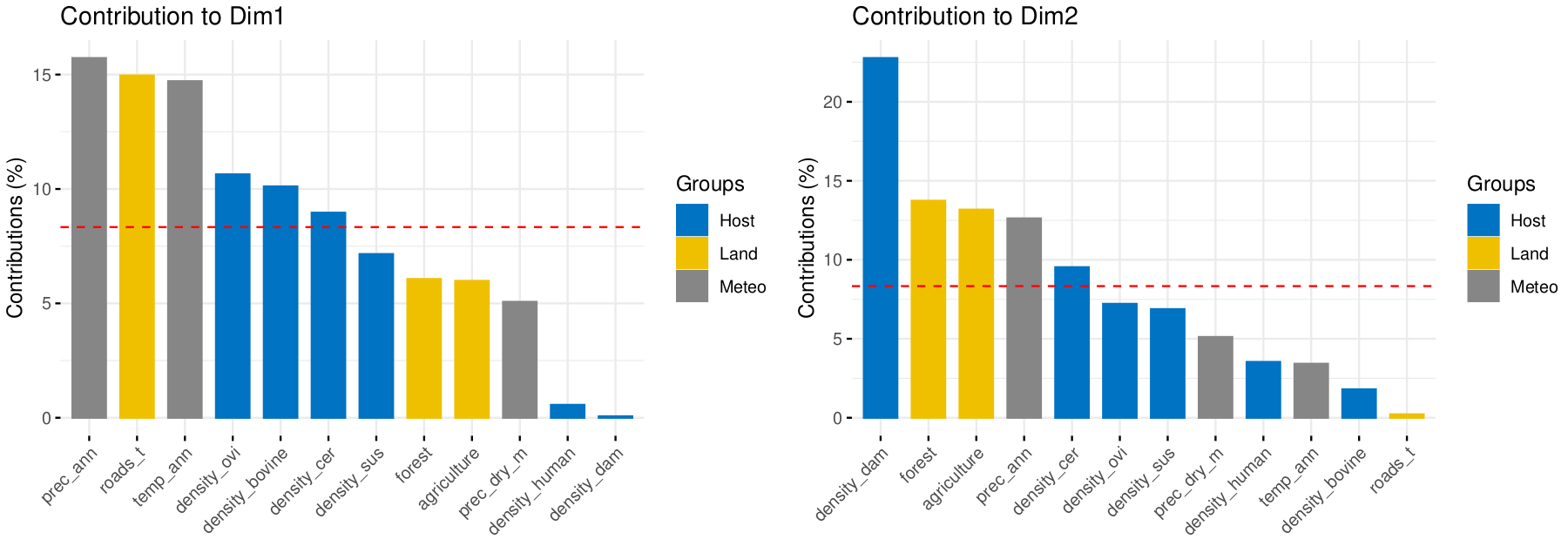

**Figure T2S6** – MFA variable contribution to dimensions 1 and 2. The red dashed line indicates the expected average value for contribution if the variables contributed uniformly (i.e. 100/N, N=number of variables).

As presented in **Figure T2S7**, we next restricted visualisation of the MFA outputs to include only these variables with the largest contributions (***temp_ann, roads_t, prec_ann, density_ovi, density_bovine, density_cer, density_dam, forest, agriculture***; N=9), while at the same time, visualizing each sample and corresponding cluster (from considering the four clustering exercise shown in **Figures T2S1-T2S4**) in MFA space.

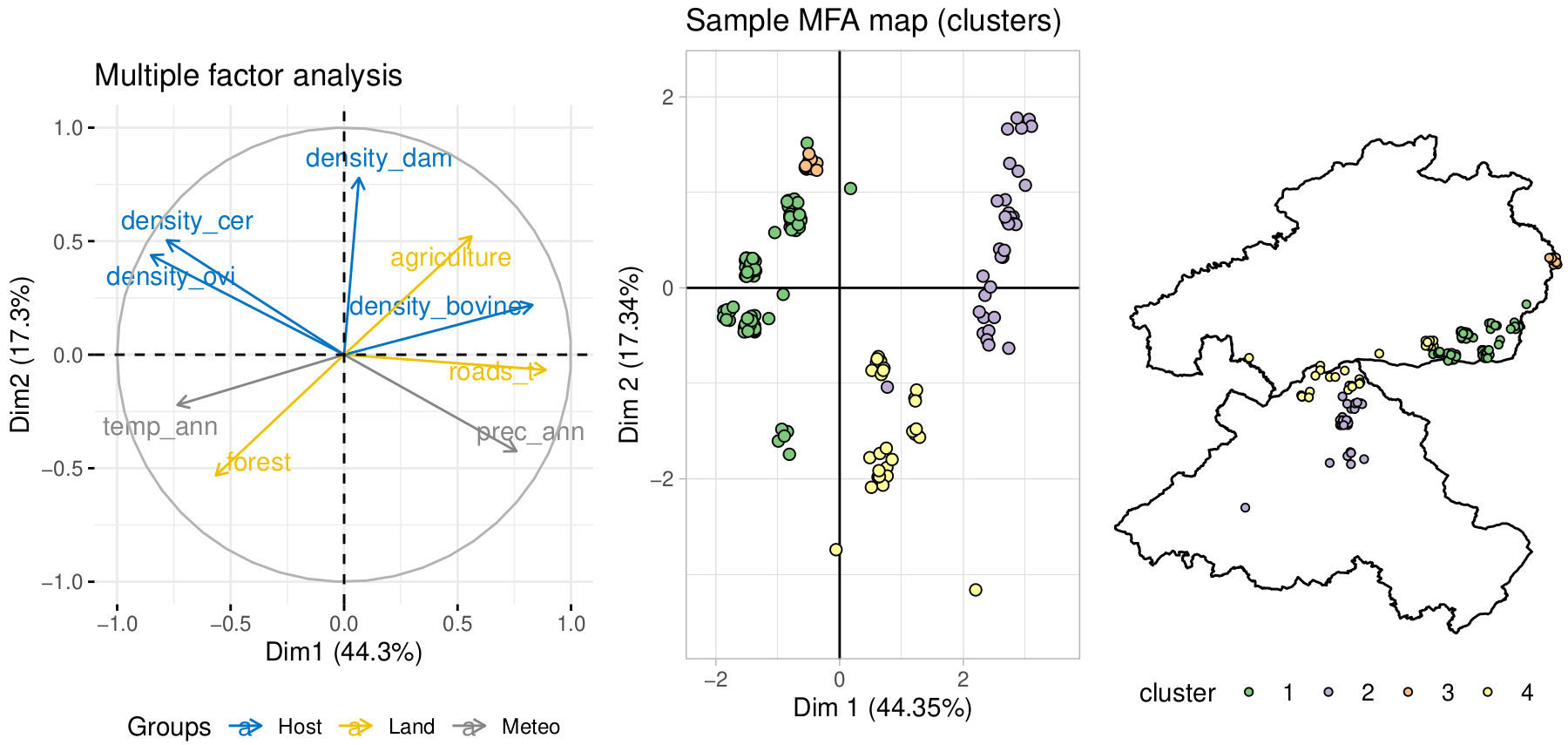

**Figure T2S7** – Sample clusters and MFA top variable visualization.

Visual inspection allowed to gain insights into inherent correlative data structures and signatures relative to each of the 4 clusters:

- Cluster 1 was characterized by a combination of high mouflon (**density_ovi**) and red deer (**density_cer**) densities and annual temperature (**temp_ann**), existing also on a negative gradient between forest and agriculture.
- Cluster 2 was composed of samples with a combination of high agriculture, low forest, high bovine density (**density_bovine**) and road density (**road_t**), and low temperature (**temp_ann**).
- Cluster 3 had high density of red deer (**density_cer**), notably being a subgroup of cluster 1 due to low mean annual temperature (**temp_ann**).
- Cluster 4 samples had a particular combination of high forest, low Fallow deer density (**density_dam**) and high precipitation (**prec_ann**).

These insights could be confirmed by visual inspection of variable gradients across the clusters as shown in **Figure T2S8**.

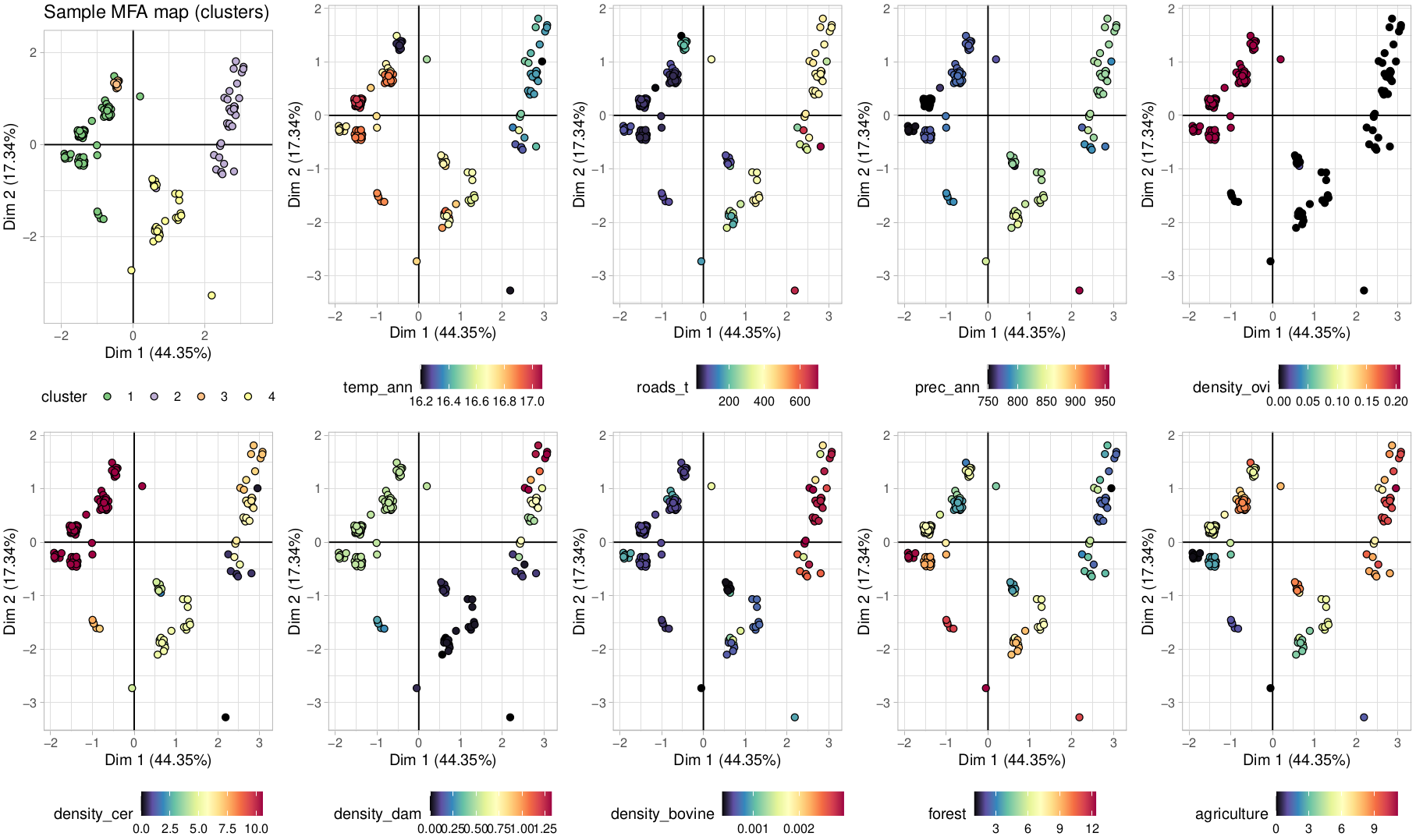

**Figure T2S8** – Sample clusters in MFA space (top contributing variables) colored by variable variation.
